# Tubulinopathy mutations in *TUBA1A* that disrupt neuronal morphogenesis and migration override XMAP215/Stu2 regulation of microtubule dynamics

**DOI:** 10.1101/2021.12.06.471490

**Authors:** Katelyn J. Hoff, Jayne E. Aiken, Mark A. Gutierrez, Santos J. Franco, Jeffrey K. Moore

## Abstract

Heterozygous, missense mutations in α- or β-tubulin genes are associated with a wide range of human brain malformations, known as tubulinopathies. We seek to understand whether a mutation’s impact at the molecular and cellular levels scale with the severity of brain malformation. Here we focus on two mutations at the valine 409 residue of TUBA1A, V409I and V409A, identified in patients with pachygyria or lissencephaly, respectively. We find that ectopic expression of *TUBA1A*-V409I/A mutants disrupt neuronal migration in mice and promote excessive neurite branching and delayed retraction in primary neuronal cultures, accompanied by increased microtubule acetylation. To determine the molecular mechanisms, we modeled the V409I/A mutants in budding yeast and found that they promote intrinsically faster microtubule polymerization rates in cells and in reconstitution experiments with purified tubulin. In addition, V409I/A mutants decrease the recruitment of XMAP215/Stu2 to plus ends and ablate tubulin binding to TOG domains. In each assay tested, the *TUBA1A*-V409I mutant exhibits an intermediate phenotype between wild type and the more severe *TUBA1A*-V409A, reflecting the severity observed in brain malformations. Together, our data support a model in which the V409I/A mutations may limit tubulin conformational states and thereby disrupt microtubule regulation during neuronal morphogenesis and migration.

## INTRODUCTION

Brain development requires cargo transport, force generation, and structural reinforcement by the microtubule cytoskeleton. The regulation of microtubules must be finely tuned to meet specific demands for different cell types, timepoints in development, and even different locations within a cell. Particularly in neurons, the spatial and temporal regulation of microtubules establishes cargo transport networks, facilitates efficient signal transduction, and supports the extension and retraction of neurites (Dent and Baas, 2014; Dent and Kalil, 2001; He et al., 2002; Lin et al., 2012; Witte et al., 2008). Accordingly, defects in microtubule regulation in neurons have been linked to brain malformations such as lissencephaly, microcephaly, and autism spectrum disorders, among others (Bahi-Buisson et al., 2014; Chakraborti et al., 2016; Srivastava and Schwartz, 2014).

Brain malformations such as lissencephaly, pachygyria, and polymicrogyria are associated with defects in neuronal migration. During neuronal migration, neurons that are born at the neuroepithelium must migrate out of the ventricular zone along radial glia cells to reach their destination at the cortical plate (Barkovich et al., 2012, 2005). Throughout the course of radial migration cortical neurons must transition through various morphologies, first by transitioning from a bipolar to a multipolar state, then back to a bipolar state to complete migration (Nadarajah et al., 2001; Noctor et al., 2004; Tabata and Nakajima, 2003). After arriving at the cortical plate, the neurons are correctly positioned and polarized to connect to each other and send and receive information. During this dynamic process of neuronal morphogenesis, microtubules generate force and provide structural support during neuronal morphogenesis. Therefore, microtubule networks must be acutely regulated during these transitions (Chesta et al., 2014; Dehmelt et al., 2006; Dent et al., 2011; He et al., 2002; Lu et al., 2013; Sainath and Gallo, 2015; Winding et al., 2016). For example, the microtubule motor, kinesin-6, is important for establishing a leading process that is required for the transition from a multipolar to bipolar state and subsequent neuronal migration (Falnikar et al., 2013). Together this suggests the critical importance of regulating microtubules in the right place and at the right time during the morphological transitions neurons must undergo for migration and, ultimately, proper brain development.

‘Tubulinopathies’ encompass a wide range of heterozygous, missense mutations in the genes encoding α- and β-tubulins that are associated with a spectrum of brain malformations. (Bahi-Buisson et al., 2014; Fallet-Bianco et al., 2014). α- and β-tubulins are encoded by a number of different genes, known as isotypes, that differ in amino acid sequence. The expression of different isotypes varies across different cell types as well as during different stages of development. Tubulinopathy mutations have been identified in three of the eight β-tubulin isotypes, and in one of the seven α-tubulin isotypes, a gene known as *TUBA1A* (Bahi-Buisson et al., 2014; Ludueña and Banerjee, 2008). *TUBA1A* is the most highly expressed α-tubulin isotype in post-mitotic neurons in the developing brain (Buscaglia et al., 2020b; Gloster et al., 1994, 1999). To date, a total of 121 heterozygous, missense mutations have been identified in *TUBA1A* and are associated with neurodevelopment disorders (Hebebrand et al., 2019). It remains unclear whether the different severities of malformations observed in these patients are a result of differences in genetic background or are a result of a specific functional difference in the mutant tubulin. A few of these *TUBA1A* mutants have been identified as loss-of-function mutants that are unable to properly assemble into microtubule polymer and thus result in an undersupply of tubulin in the cell (Belvindrah et al., 2017; Keays et al., 2007). However, other *TUBA1A* mutants appear to be gain-of-function, as they are capable of microtubule assembly and act dominantly to perturb microtubule function in migrating neurons (Aiken et al., 2020, 2019). If haploinsufficiency does not explain all *TUBA1A* tubulinopathies, then it is necessary to define clear mechanistic models for how different mutations in α-tubulin can result in different developmental outcomes. Only a small fraction of *TUBA1A* mutants have been studied, and more continued to be identified in the clinic. Therefore, it is important to establish clear mechanistic themes that will be useful in predicting the molecular, cellular, and tissue-level phenotypes of specific mutations.

The fundamental role of α-tubulin is to complex with β-tubulin into heterodimers and form microtubule polymers that are dynamic and highly regulated via both intrinsic and extrinsic factors (Bodakuntla et al., 2019; Borys et al., 2020; Goodson and Jonasson, 2018; Manka and Moores, 2018; Mitchison and Kirschner, 1984). Purified tubulin assembles into dynamic microtubules in the absence of extrinsic microtubule-associated proteins (MAPs) that are found in cells (Horio and Hotani, 1986; Mitchison and Kirschner, 1984; Walker et al., 1988). This intrinsic activity is in part regulated by the series of conformational states that tubulin undergoes as it transitions from free heterodimer into microtubule polymer (Chrétien et al., 1995; Mandelkow et al., 1991). In recent years, studies using cryo-electron microscopy and recombinant tubulin approaches have advanced the field’s understanding of the transitions that accompany GTP hydrolysis as free tubulin assembles into microtubule lattice (Geyer et al., 2018; Manka and Moores, 2018; Roostalu et al., 2020; Zheng et al., 1995). In addition to nucleotide-dependent conformational changes, tubulin must transition from a ‘curved’ free heterodimer to a straight conformation in the microtubule lattice (Buey et al., 2006; Jánosi et al., 1998; Nawrotek et al., 2011; Nogales and Wang, 2006; Ravelli et al., 2004; Rice et al., 2008). Tubulin domains that control the curved-to-straight transition are not completely understood. The ability of tubulin to adopt a range of conformational states not only impacts its intrinsic microtubule dynamics, but also tubulin’s interactions with a wide spectrum of MAPs. For example, doublecortin is predominantly expressed in the developing brain and its N-terminal doublecortin-like (DC) repeat domain binds at the corner of four tubulin subunits in a straight microtubule lattice (Fourniol et al., 2010; Manka and Moores, 2020; Moores et al., 2006). On the other hand, members of the XMAP215 protein family possess a variable number of TOG (tumor overexpressed gene) domains, some of which preferentially bind to curved tubulin (Ayaz et al., 2014, 2012; Slep and Vale, 2007). XMAP215 proteins use this activity to either pull curved tubulin toward the microtubule plus end for polymerization or drive lattice-bound tubulin into a curved state and promote catastrophe (Ayaz et al., 2014; Brouhard et al., 2008; Farmer et al., 2021; Geyer et al., 2018). Therefore, understanding the effects of tubulin conformations on both intrinsic activity and extrinsic regulatory mechanisms are essential for elucidating the regulation of microtubule dynamics.

In this study we investigate the molecular mechanism of two previously identified tubulinopathy-associated mutations at the valine 409 residue of *TUBA1A*, V409I and V409A. The patient harboring the V409A mutation exhibited lethal lissencephaly, whereas the patient with the V409I mutation presents with a milder malformation known as pachygyria (Bahi-Buisson et al., 2014; Fallet-Bianco et al., 2014). Studying these two mutations that occur at the same residue, yet are associated with varying degrees of brain malformations in patients, has potential to provide insight into how perturbing tubulin activity and the microtubule cytoskeleton in slightly different ways ultimately impacts brain development. We find that the expression of V409I or V409A dominantly disrupts neuron migration in mice *in vivo* and neuron morphologies during *in vitro* development. By modeling the V409I/A mutants at the corresponding residue in budding yeast, V410, we find that the mutants increase microtubule polymerization and depolymerization rates in cells and disrupt the regulation typically conferred by XMAP215/Stu2. We propose that these results are reconciled by the V410I/A mutants adopting a persistently straightened state as compared to WT tubulin, and this drives intrinsically faster microtubule polymerization rates *in vitro*. Importantly, we find that the severity of the V409I and V409A mutations scale from protein activity, to microtubule dynamics in cells, to neuron morphology and migration. Our results suggest that the *TUBA1A*-V409I/A mutations cause brain malformations by limiting the conformational transitions of tubulin heterodimers.

## RESULTS

### *TUBA1A*-V409I/A mutants dominantly disrupt cortical neuron migration

To determine whether *TUBA1A*-V409I/A variants act as loss- or gain-of-function mutations we first sought to establish whether the mutant proteins can polymerize into microtubules. We expressed a hexahistidine (6X-His)-tagged *TUBA1A* construct in cortical neuron cultures harvested from P0 rats. The 6X-His tag was inserted between residues I42 and G43 in a flexible loop of *TUBA1A* that allows for the addition of amino acids without perturbing tubulin function (Buscaglia et al., 2020a). After two days in culture (DIV2) we extracted soluble tubulin from the neurons, fixed the cells, and stained for the 6X-His tag. Both the *TUBA1A*-V409I and *-*V409A mutants incorporate into microtubules, similar to wild-type (WT) controls, as indicated by the microtubule filaments that can be resolved in the soma (Figure 1A). As a positive control, we expressed the *TUBA1A*-T349E mutant that does not assemble into polymer, as evidenced by the lack of labeled polymer in the soma (Figure 1A). We find that the amount of 6X-His signal retained after soluble protein extraction is similar between WT and V409I/A expressing cells (Figure 1B). These results suggest that the *TUBA1A*-V409I/A mutants assemble into microtubule polymer and have similar levels of microtubule polymer compared to WT.

**Figure 1.**
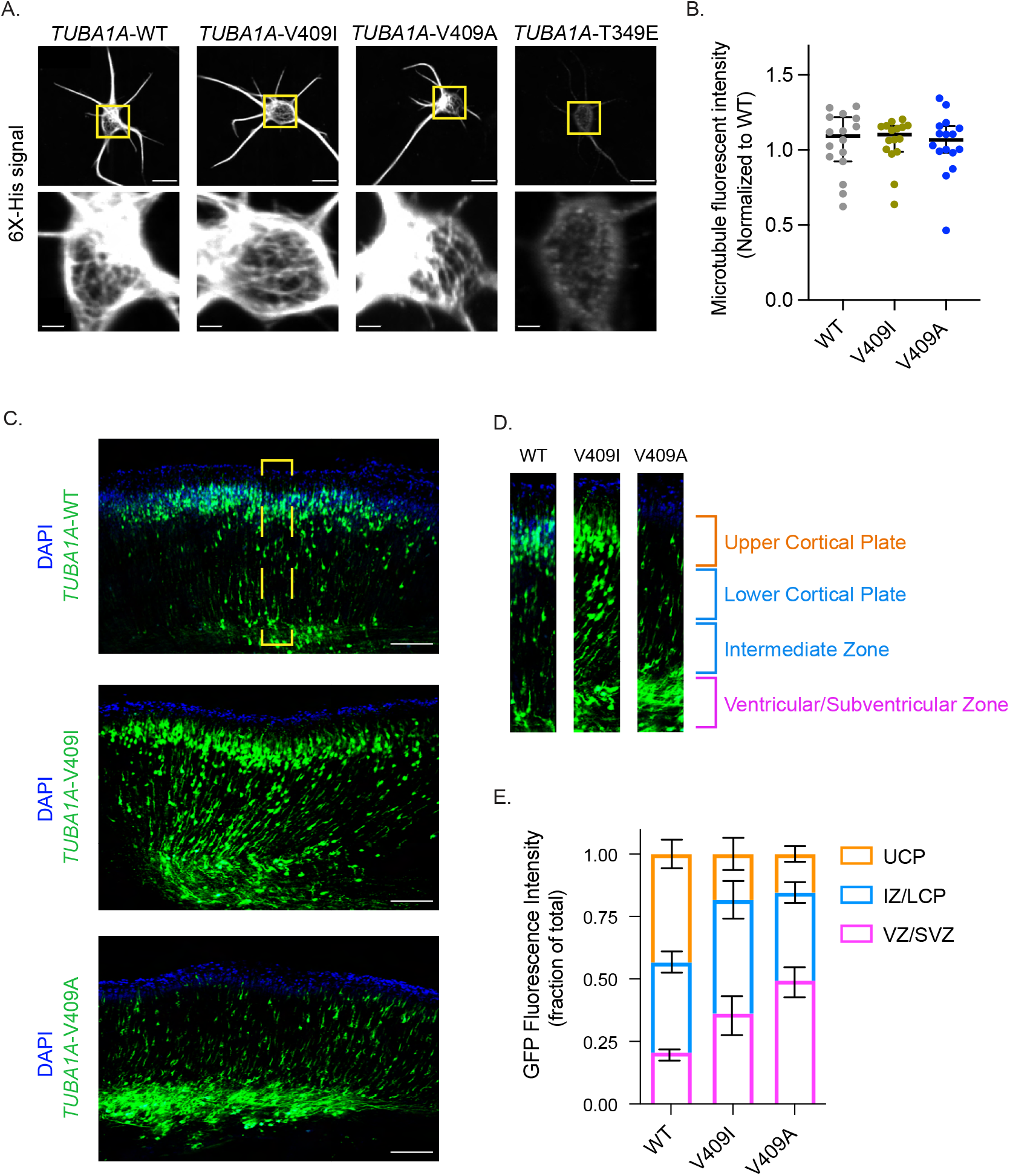
*TUBA1A*-V409I/A mutants dominantly disrupt cortical neuron migration. **A)** Representative images of neurons ectopically expressing 6X-His tagged *TUBA1A*-WT, -V409I, -V409A, and -T349E. The soluble tubulin was extracted from the cells, and then cells were stained with an anti-6X-His antibody. Bottom insets are representative of yellow boxes. Scale bar = 10μm in top panels. Scale bar = 2μm in bottom panels. At least 16 neurons were imaged for each condition. **B)** 6X-His fluorescence intensity was measured for each neuron imaged in panel A. Each dot represents a single cell. Bars are median ± 95% confidence interval. **C)** Representative coronal sections of E18.5 mouse brains that were electroporated with the above constructs at E14.5. GFP labels electroporated cells. Scale bar = 200μm. At least 18 brains were imaged for each condition. Yellow box represents region in **(D)**. **(E)** Representative cortical region was divided into 4 equal quartiles and GFP fluorescence intensity was measured in each. 1_st_ quartile = ventricular/subventricular zones (VZ/SVZ), 2_nd_ quartile = intermediate zone (IZ), 3^rd^ quartile = lower cortical plate (LCP), 4_th_ quartile = upper cortical plate (UCP). Bars are mean ± 95% confidence interval.

We next asked whether incorporation of the *TUBA1A*-V409I/A mutants into microtubules *in vivo* is sufficient to disrupt cortical neuron migration. To answer this question, we performed *in utero* electroporations to ectopically express *TUBA1A* in migrating neurons in the developing mouse brain. At embryonic day (E) 14.5, we electroporated neural progenitors in the cortical ventricular zone, which give rise to immature excitatory neurons that migrate from the ventricular zone into the upper cortical plate. The cDNA expression plasmids used for the electroporations encode for either WT or mutant *TUBA1A*, along with GFP to identify the electroporated cells. The majority of GFP+ neurons ectopically expressing *TUBA1A*-WT successfully migrate from the ventricular zone to the upper cortical plate by E18.5 (Figure 1C). In contrast, neurons ectopically expressing *TUBA1A*-V409I do not migrate to the upper cortical plate as robustly as WT control cells by E18.5, as evidenced by many neurons remaining in the ventricular, subventricular, and intermediate zones. Strikingly, a majority of *TUBA1A*-V409A expressing neurons fail to migrate at all by E18.5 and remain primarily in the ventricular and subventricular zones. To quantify migration for comparison across experiments, we divided the cortical sections into four equal segments from the ventricular zone to the top of the cortical plate and measured the proportion of GFP signal in each segment (Figure 1D). Approximately, the first quartile represents the ventricular and subventricular zones, the second quartile represents the intermediate zone, the third quartile represents the lower cortical plate, and the fourth quartile represents the upper cortical plate. We find that the largest proportion of *TUBA1A*-WT expressing cells (43.4%) are found in the upper cortical plate region (labeled in orange), while the largest proportion of *TUBA1A*-V409I expressing cells (45.4%) are in the intermediate zone and lower cortical plate, and the largest proportion of *TUBA1A*-V409A expressing cells (49.4%) are in the ventricular/subventricular zones (Figure 1E). In each of our experiments, untransfected control cells successfully reach the cortical plate, as evidenced by abundant DAPI signal in the upper cortical plate segment, indicating that the radial glia cells that support migration are not impaired by the transfection of mutant *TUBA1A* in our experiments. These results suggest that the *TUBA1A*-V409I/A mutants act dominantly to disrupt neuron migration, with *TUBA1A*-V409A being more severe than the *TUBA1A*-V409I mutant.

### *TUBA1A*-V409I/A mutants increase neurite branching and delay neurite retraction

The microtubule cytoskeleton plays a crucial role in the series of morphological changes that cortical neurons must undergo throughout the course of radial migration (Nadarajah et al., 2001; Noctor et al., 2004; Tabata and Nakajima, 2003). When immature neurons are born, they extend numerous neurites to probe their environment for directional cues. Once these cues have been identified, neurons must retract most of their neurites and become bipolar, such that one neurite becomes the axon and a neurite on the opposite side of the cell becomes the leading process that guides radial migration. Therefore, it is particularly crucial that microtubules are able to deftly respond to various cues throughout development that promote these different morphology transitions. Therefore, we sought to determine if the *TUBA1A*-V409I/A mutant microtubules perturb neuron morphologies during *in vitro* development. We formed two distinct hypotheses of how *TUBA1A*-V409I/A mutant microtubules could affect neuron morphogenesis: 1) insufficient neurite extension, or 2) insufficient neurite retraction.

To distinguish between these two hypotheses, we first compared the extension of neurites between WT and V409I/A mutants by ectopically expressing the 6X-His-tagged *TUBA1A* (WT or mutant) plasmids described above. We then quantified the number of primary, secondary, and tertiary neurites at two morphologically distinct stages of *in vitro* cortical neuron development, stage 2 (DIV1) and stage 3 (DIV2). Stage 2 neurons are characterized as being multipolar with all neurites being approximately similar in length (Dotti et al., 1988). On average, there is no appreciable difference in the number of neurites quantified in stage 2 cells expressing WT or V409I/A mutant *TUBA1A*. Most neurons at stage 2 have four primary neurites, and we rarely observe secondary or tertiary branches (Supplemental Figure 1A and B). Consistent with this finding, stage 3 WT and V409I/A mutant neurons have a similar number of primary neurites (5.123 ± 0.426, 4.911 ± 0.366, 5.356 ± 0.404 for WT, V409I, and V409A, respectively; Figure 2A and B). However, *TUBA1A*-V409I expressing cells exhibit a slight, but measurable increase in secondary branches as compared to WT cells (0.600 ± 0.227, 0.893 ± 0.264 branches/cell for WT and V409I, respectively; Figure 2B). Strikingly, *TUBA1A*-V409A expressing cells have a significantly greater number of secondary branches, and some also exhibit tertiary branches (Secondary: 0.600 ± 0.227, 2.186 ± 0.682 branches/cell, Tertiary: 0.015 ± 0.031, 0.322 ± 0.171 branches/cell for WT and V409A, respectively; Figure 2B). These data fail to support the hypothesis that *TUBA1A*-V409I/A mutants result in insufficient neurite extension because we find a similar number of primary neurites in WT and mutant expressing cells. Our data instead lend support to the alternative hypothesis that there is insufficient neurite retraction, as evidenced by the increase in neurite branching.

**Figure 2.**
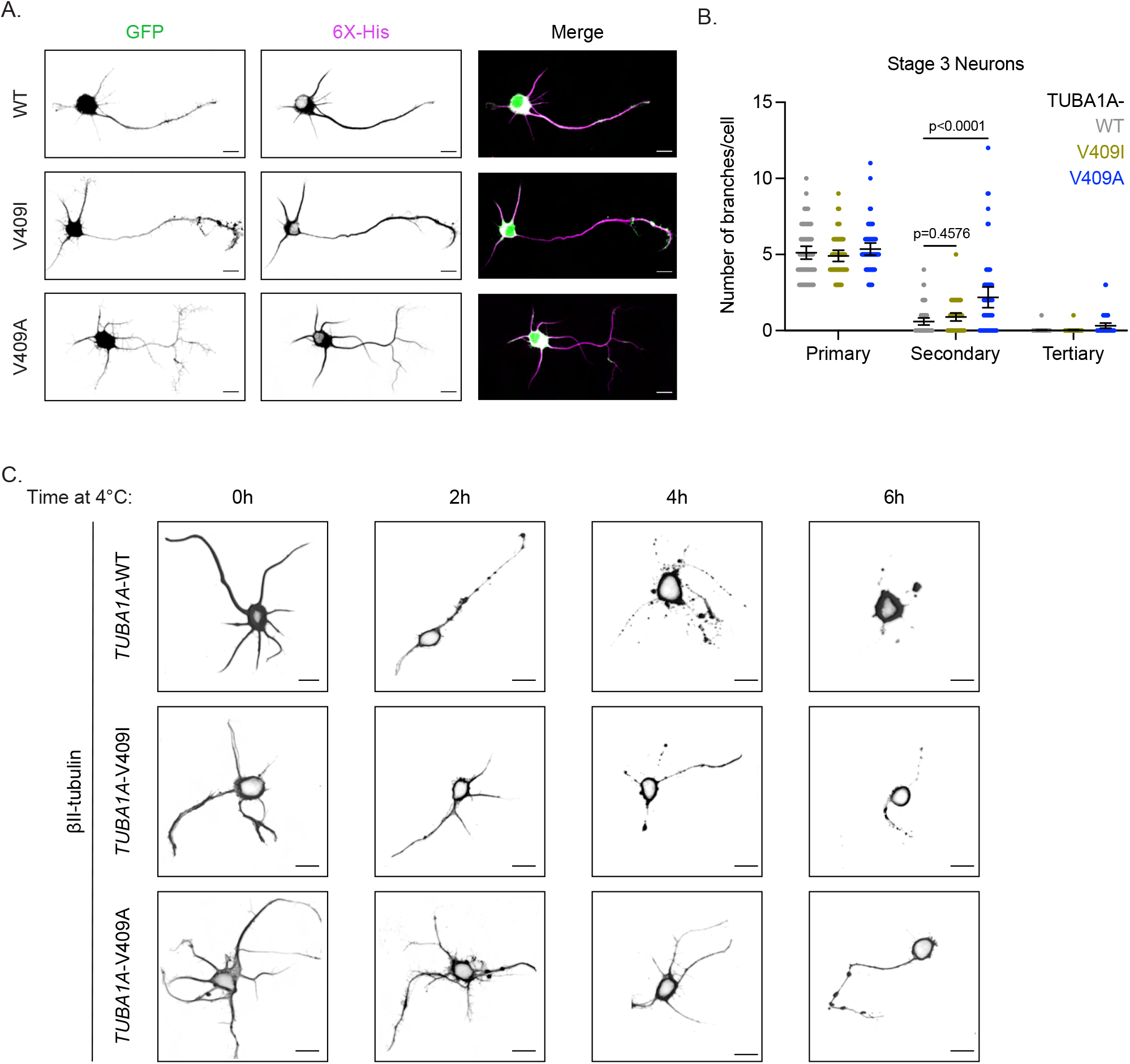
*TUBA1A*-V409I/A mutants alter neuron morphologies. **A)** Representative DIV2 cortical neurons expressing cytoplasmic GFP and 6X-His tagged *TUBA1A*-WT, -V409I, and -V409A. Scale bar = 10μm. At least 27 neurons were imaged for each condition. **B)** Quantification of number of primary, secondary, and tertiary branches in each condition. Each dot represents a single cell. Bars are mean ± 95% confidence interval. **C)** Representative images of neurons expressing the above plasmids, exposed to 4°C for indicated time, and stained with TUBB2A/B. Scale bar = 10μm.

To address this, we tested whether the neuron microtubules respond to destabilizing cues that are necessary for neurite retraction. We stimulated neurite retraction *in vitro* by placing the neuron cultures at 4°C over the course of six hours. At low temperatures, microtubules destabilize, which ultimately leads to neurite retraction (Breton and Brown, 1998; Tilney and Porter, 1967; Weber et al., 1975). During our time course experiment, samples from each genotype were fixed at two-hour intervals and stained for β-II tubulin, a highly abundant β-tubulin isotype in cortical neurons, to label microtubules in neurites. After two hours at 4°C, *TUBA1A*-WT expressing cells (identified by cytoplasmic GFP) lost most neurites, and those that remain formed the retraction bulbs that are characteristic of retracting neurites (Figure 2C and Supplemental Figure 1C). By four and six hours of cold exposure, β-II tubulin signal is only found in the cell soma of neurons expressing *TUBA1A*-WT. In contrast, most *TUBA1A*-V409I expressing cells do not exhibit retraction bulbs or neurite loss until four hours of 4°C exposure. The most severe defect is detected in *TUBA1A*-V409A expressing cells where neurites are maintained throughout most of the time course, with retraction bulbs beginning to form only after six hours of the 4°C exposure. Together, these data suggest that expression of *TUBA1A*-V409I or -V409A creates hyper-stable neurites and, particularly in the case of V409A, excessive neurite branching.

### *TUBA1A*-V409I/A mutant microtubules have high levels of acetylation

We next sought to determine if the increase in neurite branching and resistance to cold-induced neurite retraction seen in *TUBA1A*-V409I/A expressing cells is a result of altered microtubule dynamics. To test this, we quantified levels of tubulin post-translational modifications in WT- and mutant-expressing cells. Tyrosinated tubulin is a marker of dynamic, newly formed microtubules and is typically evenly distributed amongst the axon and dendrites (reviewed in Westermann and Weber, 2003). Acetylated tubulin is a marker of stable microtubules and is highly localized to the axon, but largely absent in the dendrites.

As expected, stage 3 control cells ectopically expressing *TUBA1A*-WT have acetylated tubulin localized to the axon and tyrosinated tubulin throughout the neuron (Figure 3A and B). Both *TUBA1A*-V409I and -V409A expressing neurons also exhibit tyrosinated tubulin throughout the neuron, similar to that observed in control cells expressing *TUBA1A*-WT (Axons: 0.487 ± 0.061, 0.507 ± 0.069, 0.471 ± 0.070A.U. and Dendrites: 0.524 ± 0.069, 0.511 ± 0.072, 0.543 ± 0.083A.U. for WT, V409I, and V409A, respectively; Figure 3A and B). *TUBA1A*-V409I expressing cells have a modest increase in acetylated tubulin in both the axon and dendrites, and *TUBA1A*-V409A expressing cells exhibit a striking increase in microtubule acetylation in all neurites and even in the soma (Axons: 2.861 ± 0.227, 3.025 ± 0.253, 3.721 ± 0.295A.U. and Dendrites: 1.239 ± 0.204, 1.492 ± 0.145, 2.395 ± 0.184A.U. for WT, V409I, and V409A, respectively; Figure 3A and C). Although acetylated tubulin levels are increased in the *TUBA1A*-V409I/A mutants, our experiments where soluble tubulin is extracted indicate that the total amount of microtubule polymer is similar between WT and mutant-expressing cells (Figure 1B). This suggests that increased microtubule acetylation in cells expressing V409I/A mutants is not simply attributable to increased microtubule polymer, but rather increased levels of acetylation on the microtubules.

**Figure 3.**
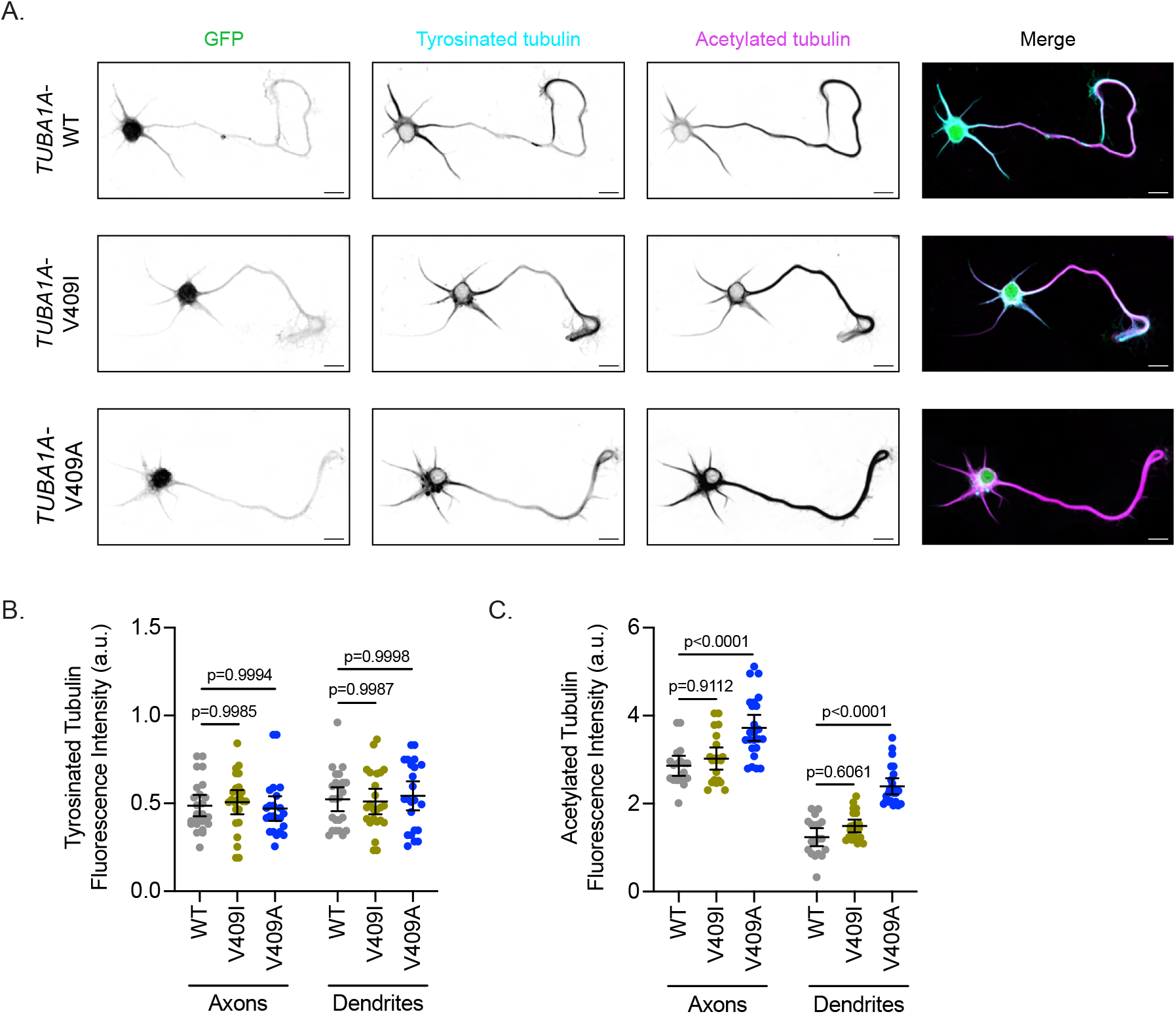
*TUBA1A*-V409I/A microtubules have increased acetylation levels. **A)** Representative DIV2 cortical neurons expressing plasmids with cytoplasmic GFP and 6X-His tagged *TUBA1A*-WT, -V409I, and -V409A. Cells stained with tyrosinated tubulin and acetylated tubulin. Scalebar = 10μm. Quantification of tyrosinated tubulin **(B)** and acetylated tubulin **(C)** fluorescence intensity in a cell’s axon and dendrites. Each dot represents quantification of a single cell. Bars are mean ± 95% confidence interval.

### Modeling *TUBA1A*-V409I/A mutants in budding yeast reveals altered microtubule dynamics

The α-tubulin V409 residue is highly conserved across eukaryotes (Figure 4A). To better understand the molecular impact of the *TUBA1A*-V409I/A mutants, we created analogous mutants at the corresponding residue in *Saccharomyces cerevisiae* (or budding yeast), V410. Using this system, we created strains in which all the α-tubulin expressed in the cell was either WT, V410I, or V410A. We find that compared to WT, V410I cells have no significant change in fitness, while V410A cells exhibit a slight fitness defect, as indicated by a 4.4% increase in doubling time (Supplemental Figure 2A). By introducing either the V410I or V410A mutation into both α-tubulin isotypes of budding yeast (*TUB1* and *TUB3*), we can measure the dynamics of individual microtubules that consist of a homogenous supply of either WT or mutant α-tubulin. Thus, any effect we see on dynamics would be a result of the mutant of interest as opposed to a compensatory response by alternative tubulin isotypes.

**Figure 4.**
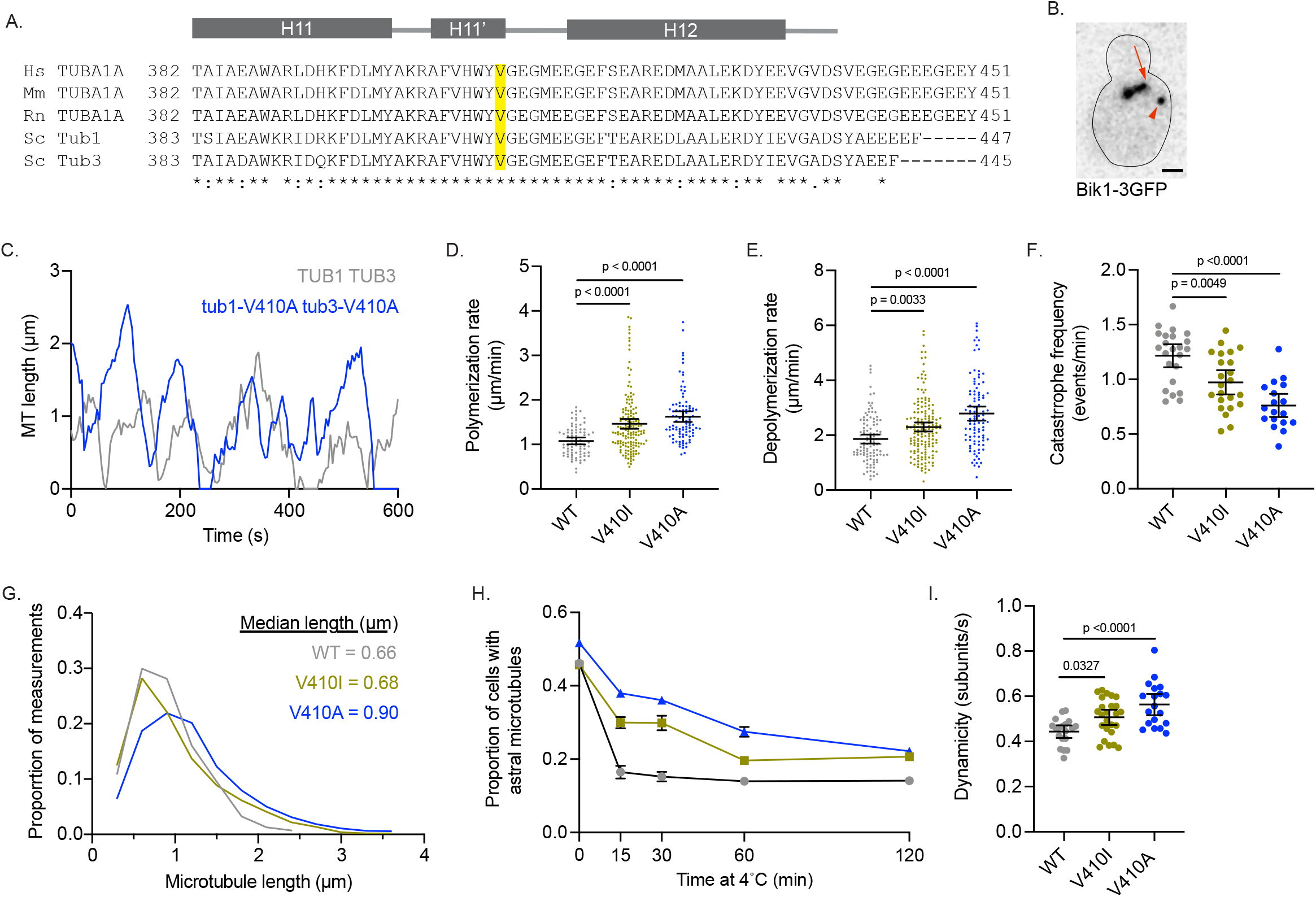
Modeling *TUBA1A*-V409I/A mutants in budding yeast reveals altered microtubule dynamics. **A)** Sequence alignment of human, mouse, and rat TUBA1A and budding yeast α-tubulins Tub1 and Tub3. Valine 409 is highlighted in yellow and resides in a highly conserved helix known as H11’. **B)** Representative image of budding yeast cell with microtubule plus-end binding protein Bik1 labeled with 3GFP. Scalebar = 1μm. **C)** Life plot of the dynamics of individual microtubules from WT (gray) and V410A (blue) cells showing the change in length over time. **D)** Polymerization rates of astral microtubules. Each dot represents a single polymerization event. Bars are mean ± 95% confidence interval. **E)** Depolymerization rates of astral microtubules. Each dot represents a single depolymerization event. **F)** Catastrophe frequency for astral microtubules expressed as the number of catastrophe events that occur per minute. Each dot represents the average catastrophe frequency of a single cell. **G)** Histogram of all astral microtubule lengths from time lapse imaging of WT, V410I, and V410A cells. **H)** Proportion of cells with astral microtubules after indicated time at 4°C. Symbols represent average of 3 independent experiments where at least 295 cells were analyzed per condition. **I)** Dynamicity of astral microtubules calculated as the total change in length divided by the total change in time. Each dot represents the calculated dynamicity value of a single cell. For all measured dynamics, at least 28 cells were measured for each genotype.

To track the length of individual microtubules over time in cells, we used Bik1-3GFP, the yeast homologue of CLIP-170, as a marker for microtubule plus-ends (Figure 4B and 4C). Compared to WT, we find that *tub1*/*tub3*-V410A mutant microtubules have significantly faster polymerization rates, while *tub1*/*tub3*-V410I microtubules have intermediate rates between WT and V410A mutants (1.078 ± 0.078, 1.463 ± 0.112, 1.625 ± 0.117µm/min for WT, V410I, and V410A, respectively; Figure 4D). Similarly, *tub1*/*tub3*-V410A microtubules, and to a lesser extent -V410I microtubules, have faster depolymerization rates compared to WT (1.861 ± 0.165, 2.300 ± 0.166, 2.788 ± 0.256µm/min for WT, V410I, V410A, respectively; Figure 4E). Our data also shows that *tub1*/*tub3*-V410A microtubules exhibit very few catastrophes compared to WT, and the *tub1*/*tub3*-V410I microtubules again have an intermediate catastrophe frequency (1.216 ± 0.105, 0.973 ± 0.111, 0.761 ± 0.106 events/minute for WT, V410I, and V410A, respectively; Figure 4F). Accordingly, *tub1*/*tub3*-V410A mutant microtubules reach longer median lengths than either the *tub1*/*tub3*-V410I or WT microtubules (0.66, 0.68, 0.90µm for WT, V410I, V410A, respectively; Figure 4G). Additionally, we find that astral microtubules in *tub1/tub3*-V410A mutant cells are retained for a longer time at 4°C than those in WT control cells, and *tub1/tub3*-V410I microtubules have an intermediate phenotype (Figure 4H). Summarizing these dynamics data, we find that α-tubulin-V410I/A microtubules, and particularly the *tub1*/*tub3*-V410A variant, exhibit faster microtubule polymerization rates and decrease how often the microtubule catastrophes. However, when these mutant microtubules do catastrophe, they depolymerize at a faster rate than WT.

To understand how the different microtubule parameters described above work together to influence microtubule activity, we calculated the dynamicity of WT and *tub1*/*tub3*-V410I/A microtubules. Dynamicity is defined as the total change in microtubule length divided by the change in time (Jordan et al., 1993). We find that *tub1*/*tub3*-V410A microtubules have the highest dynamicity values, while *tub1*/*tub3*-V410I microtubules have an intermediate dynamicity between -V410A and WT (0.443 ± 0.028, 0.507 ± 0.034, 0.563 ± 0.047 subunits/second; Figure 4I). Similar to our work in neurons, the V410A mutant has the strongest effect on microtubule dynamics while the V410I mutant has a more intermediate impact. Together, the results of modeling these patient-associated mutations in budding yeast indicates that the α-tubulin-V410I/A mutations are sufficient to alter microtubule dynamics in this system.

### *tub1/tub3*-V410I/A microtubules have decreased localization of and affinity for XMAP215/Stu2

Microtubule dynamics in cells are the product of both intrinsic tubulin activity, as well as regulation by extrinsic MAPs. Therefore, we sought to determine how *tub1/tub3*-V410I/A affect extrinsic and/or intrinsic modes of regulation to alter microtubule dynamics. The α-tubulin V409 (human) or analogous V410 (yeast) residue resides on the external surface of the tubulin heterodimer near the binding sites of a wide variety of microtubule-associated proteins (Löwe et al., 2001). In particular, one structural analysis highlights the α-tubulin V410 residue as a potential interactor with the TOG2 domain of Crescerin1 (Das et al., 2015). The Crescerin1 TOG2 domain has a similar structure to the TOG1 domain of Stu2, the yeast homologue of XMAP215. As Stu2/XMAP215 is the major microtubule polymerase in cells, we predicted that the increase in microtubule polymerization observed in *tub1/tub3*-V410I/A cells could be a result of increased Stu2 activity on the mutant microtubules. To test this, we first used Stu2-3GFP to measure Stu2 localization at astral microtubule plus ends in cells where all the α-tubulin is either WT, V410I, or V410A mutant (Figure 5A). We find that *tub1/tub3*-V410A mutant microtubules have significantly decreased Stu2-3GFP fluorescence intensity at the plus ends compared to WT, while *tub1/tub3*-V410I microtubules have an intermediate phenotype between the two (3203 ± 333, 1860 ± 165, 1365 ± 143A.U. for WT, V410I, and V410A, respectively; Figure 5B). Thus, despite having increased microtubule polymerization rates, the *tub1/tub3*-V410I/A mutant microtubules have less Stu2 at microtubule plus ends.

**Figure 5.**
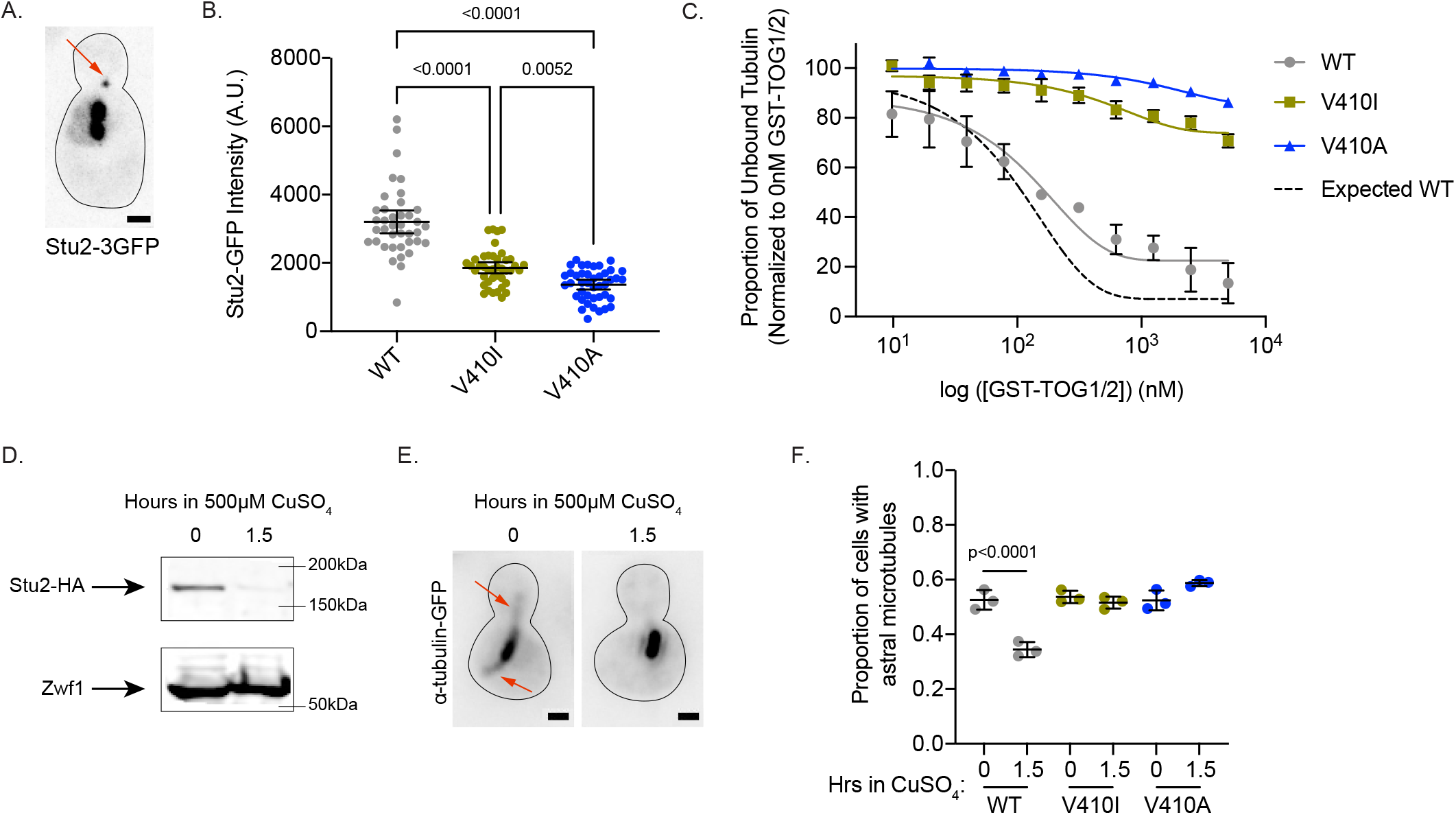
V410I/A mutants subvert Stu2 regulation. **A)** Representative image of budding yeast cell expressing Stu2-3GFP. Red arrow indicates Stu2 at microtubule plus end. Scale bar = 1μm. **B)** Quantification of Stu2-3GFP fluorescence intensity at microtubule plus ends. Each dot represents the quantification from a single cell. Bars are mean ± 95% confidence interval. At least 39 cells were analyzed per condition. **C)** Proportion of unbound tubulin in solution in the presence of increasing concentrations of GST-TOG1/2. Data was analyzed from the α-tubulin signal on supernatant samples run on western blots. Each condition was normalized to the constant amount of tubulin added to each sample in the absence of GST-TOG1/2. Dots represent averages from three separate experiments. Bars are standard error of the mean. **D)** Western blot of protein lysate from cells induced with 500μM CuSO_4_ for 0 and 1.5 hours. Blots were probed for HA and Zwf1. **E)** Example images of WT cells induced with 500μM CuSO_4_ for 0 and 1.5 hours stained for α-tubulin. Red arrows indicate astral microtubules. Scale bar = 1μm. **F)** Quantification of proportion of cells imaged that have at least one astral microtubule. Each dot represents the average proportion of cells from three separate experiments. At least 800 cells were measured for each condition. Bars are mean ± standard error of the mean.

Stu2 is comprised of two TOG domains (TOG1 and TOG2) that bind specifically to free tubulin heterodimer and are necessary for concentrating tubulin at microtubule plus ends to promote microtubule polymerization. Disrupting the interaction between the Stu2 TOG2 domain and tubulin heterodimers decreases the localization of Stu2 to microtubule plus ends (Geyer et al., 2018). We therefore predicted that the decreased localization of Stu2 at *tub1/tub3*-V410I/A microtubule plus ends may be a result of lowered affinity between the mutant tubulins and Stu2 TOG domains. To test this prediction, we purified tubulin heterodimer from budding yeast in which the purified α-tubulin was either Tub1-WT, -V410I, or -V410A. Additionally, we purified the two TOG domains in Stu2 fused to a GST tag (referred to as GST-TOG1/2). Using a low concentration of either WT or V410I/A mutant tubulin in the presence of increasing concentrations of GST-TOG1/2, we find that V410I, and more significantly V410A, mutant tubulin has decreased affinity for GST-TOG1/2 as compared to WT (Figure 5C and Supplemental Figure 3A). For reference we used previously published K_D_ values for WT yeast tubulin affinity for either TOG1 or TOG2 to determine an expected binding curve (Geyer et al., 2015). These results indicate that the decrease in Stu2 localization we see at the *tub1/tub3*-V410I/A microtubule plus ends is a consequence of the decreased affinity of Stu2 TOG1/2 domains for the mutant tubulin.

Stu2 is an essential gene in budding yeast, presumably because of its important role in regulating microtubule dynamics (Wang and Huffaker, 1997). Based on the increased microtubule polymerization rates observed for V410I/A microtubules, and the decreased affinity between V410I/A and Stu2 TOG domains, we predicted that *tub1/tub3*-V410I/A microtubules would be resistant to the depletion of Stu2. To test this, we constructed *tub1*-V410I or -V410A yeast strains where we could conditionally deplete Stu2 (Kosco et al., 2001). Upon addition of copper to the media of these cells, synthesis of *STU2* mRNA is repressed and Stu2 is degraded. We find that after 1.5 hours in 500µM copper sulfate (CuSO_4_), Stu2-HA is almost undetectable via western blot (Figure 5D and Supplemental Figure 3B). Under these conditions, the proportion of WT cells with visible astral microtubules decreases from 53% to 34% (0.526 ± 0.036, 0.345 ± 0.27 at 0 and 1.5 hours, respectively; Figure 5E-F). In contrast, *tub1*-V410I and *tub1*-V410A cells have no significant decrease in proportion of cells that have astral microtubules after 1.5 hours of Stu2 depletion (V410I: 0.537 ± 0.023, 0.516 ± 0.022, V410A: 0.524 ± 0.036, 0.588 ± 0.011 at 0 and 1.5 hours, respectively; Figure 5E-F). These data indicate that V410I and V410A microtubules persist in the absence of Stu2, and therefore the mutant tubulins may not require Stu2 polymerase activity to the same extent as WT.

### Purified *tub1*-V410A has increased microtubule polymerization rates *in vitro*

Stu2 TOG domains preferentially bind to the kinked conformation of tubulin and have low affinity for straight heterodimers (Ayaz et al., 2014, 2012). Based on our *in vitro* binding assays, we hypothesized that α-tubulin V410I/A pushes the heterodimer into a straightened state. The V410 residue resides in helix 11’ of α-tubulin, which is located at the hinge point between α- and β-tubulin (Löwe et al., 2001), and makes it a prime candidate for affecting the conformational states of the heterodimer. We predicted that a straighter heterodimer would increase microtubule polymerization rates as a straight heterodimer is more compatible with forming the microtubule lattice. Therefore, we reasoned that if the changes observed in V410I/A microtubule dynamics are a result of a straightened heterodimer conformation, the mutants should have intrinsically faster polymerization rates *in vitro*.

To test this, we used interference reflection microscopy (IRM) to measure the intrinsic polymerization activity of our purified yeast Tub1-WT, -V410I, and -V410A heterodimers in an *in vitro* dynamics assay. We used GMPCPP-stabilized, rhodamine-labeled porcine microtubule ‘seeds’ attached to the surface of the coverslip via anti-rhodamine antibodies to nucleate the assembly of microtubules from the purified tubulin in the reaction (Figure 6A). We used three concentrations of purified tubulin, between 0.5 and 0.9µM, to measure microtubule dynamics *in vitro*. Within this concentration range, we find that soluble tubulin assembles from the stabilized seeds and forms dynamic microtubules. If tubulin concentration is too high, the tubulin will spontaneously nucleate away from the stabilized seeds (to form microtubules, oligomers, aggregates, etc.) and will not be visible on the microscope. If the tubulin concentration is too low, the tubulin will not sustain assembly from the seeds. In our experimental set up, we are unable to observe microtubule dynamics below 0.5µM tubulin. For each concentration, we measured the change in microtubule length over time of WT or V410I/A mutant tubulin in a completely purified system using unlabeled tubulin (Figure 6B, Supplemental Video 1).

**Figure 6.**
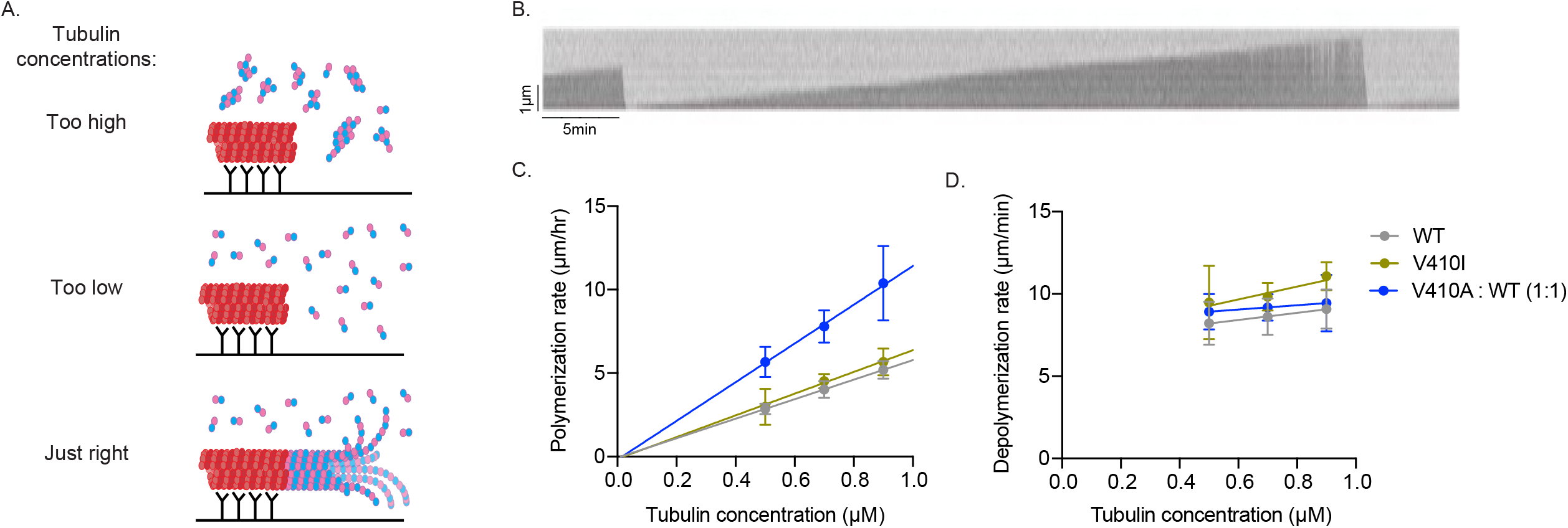
V410A has intrinsically faster microtubule polymerization rates *in vitro*. **A)** Graphic depicting aggregates and non-seeded nucleation when tubulin concentrations are too high, no nucleation when concentrations are too low, and dynamic microtubules when concentrations are at an appropriate level. **B)** Example kymograph of dynamic microtubule from purified budding yeast tubulin measured using interference reflection microscopy. X and Y scale bars = 5 minutes and 1μm, respectively. **C)** Microtubule polymerization rates at 0.5, 0.7, and 0.9μM of tubulin. Each data point represents the mean of three independent experiments in which at least 28 dynamic microtubules were measured per condition. Error bars are mean ± 95% confidence interval. **D)** Microtubule depolymerization rates.

In contrast to our results in budding yeast cells, we find that purified tub1-V410I microtubules do not have significantly increased polymerization rates as compared to WT at any of the three concentrations tested (Figure 6C). However, with purified tub1-V410A tubulin, we were unable to observe microtubule dynamics at these three concentrations, nor did we observe any at higher concentrations up to 1.5µM or lower concentrations down to 0.1µM (data not shown). Based on our previous data, we predict that tub1-V410A tubulin exhibits increased assembly activity and would be more dynamic at lower concentrations. If tub1-V410A tubulin exhibits higher assembly activity than WT tubulin, even the low end of our usable concentration range may be too high for this mutant, and we would be unable to visualize microtubule dynamics because the mutant may readily assemble away from seeds (similar to top panel of Figure 6A). To circumvent these issues while still testing the intrinsic capabilities of tub1-V410A, we mixed WT and tub1-V410A tubulin in a one-to-one ratio, for a total tubulin concentration of 0.5, 0.7, or 0.9µM. At this WT:V410A one-to-one ratio, we find an increase in microtubule polymerization rates at each concentration tested (Figure 6C). Since the amount of either WT or V410A tubulin that is present in each of these one-to-one mixtures (i.e., 0.25, 0.35, or 0.45µM) is not sufficient to support microtubule assembly on its own, we conclude that the increased microtubule polymerization is a synergistic effect of the blend of WT and V410A mutant tubulin.

The fitted lines of the data collected at each tubulin concentration provide us with important information about the intrinsic properties of the WT and mutant microtubules formed *in vitro*. The slopes of the fitted lines represent the concentration-dependent polymerization rate, and we find that WT/V410A has a ∼2-fold increased rate as compared to WT and V410I (11.6 compared to 5.8 and 6.5µm/hr/µM). The x- and y-intercepts represent the critical concentration and apparent off rate constants, respectively (x-intercepts = 0.01, 0.02, and 0.02µM; y-intercepts = -0.07, - 0.13, and -0.18µm/hr for WT, V410I, and WT/V410A, respectively). In contrast to the increase observed in microtubule polymerization rates, the microtubule depolymerization rates were not different between WT and either the V410I or V410A mutants (Figure 6D). These data indicate that *tub1*-V410A has significantly increased intrinsic microtubule polymerization as compared to WT. Together, our data explain the intrinsic mechanism by which the V409/V410A mutant has the most severe phenotypes across scales, while V409/V410I has more mild effects.

## DISCUSSION

The missense, heterozygous mutations identified in human tubulin genes and associated with brain malformations are numerous and varied (Bahi-Buisson et al., 2014; Fallet-Bianco et al., 2014). To date, we do not have a good understanding of how different mutations in tubulin genes have such a wide spectrum of detrimental effects on brain development. Investigating the molecular impact of different tubulinopathy mutations and how these mutants affect greater tissue level development is critical to advancing our understanding of the role of the microtubule cytoskeleton during neurodevelopment and disease. In this study, we aim to determine the mechanism of two different substitutions at the valine 409 residue of *TUBA1A* that are associated with different severities of brain malformations. We find that ectopically expressed *TUBA1A*-V409I/A mutants assemble into microtubules in neuron cultures and dominantly disrupt radial neuron migration, and most significantly in cells expressing *TUBA1A*-V409A (Figure 1). Our work indicates that the mutation associated with the most severe brain phenotype, *TUBA1A*-V409A, also has the most severe impact on neurite branching and microtubule dynamics, while the milder -V409I mutant has intermediate phenotypes. These findings provide evidence that the impact of tubulinopathy mutations scale up from altered microtubule polymerization activity and loss of regulation by microtubule-associated proteins to cellular and tissue defects.

As neurons migrate from the ventricular zone to the cortical plate, they must transition through different polarization states (Nadarajah et al., 2001; Noctor et al., 2004; Tabata and Nakajima, 2003). The overexpression or depletion of a wide variety of signaling molecules and cytoskeletal proteins delay or inhibit neuron morphology transitions that impair subsequent migration (reviewed in Cooper, 2014). Several cases of brain malformations similar to those investigated in this study have been linked to mutations that disrupt radial migration by inhibiting the multipolar-to-bipolar transition (e.g., loss-of-function Cdk5 mutations; Ohshima et al., 2007). While there are numerous factors that are required for proper neuron migration, our data support a hypothesis that *TUBA1A*-V409I/A mutants disrupt and/or delay proper morphology transitions. At the cellular level, we find that neurons ectopically expressing *TUBA1A*-V409I and -V409A disrupt neuron morphogenesis by promoting excessive branch stabilization (Figure 2A and B). Additionally, neurons expressing *TUBA1A*-V409I/A are more resistant to cold temperature-induced neurite retraction as compared to cells solely expressing WT tubulin (Figure 2C). To the best of our knowledge, there are no tubulinopathy mutations to date that disrupt neuron migration via impaired neuron morphology transitions. Therefore, future work will be required to determine whether this is a common mechanism for particular types of brain malformations.

Our findings support the idea that microtubule stability plays a determining role in neuron branching and morphogenesis. Previous studies have identified that taxol-treated neurons have an increase in neurite extension, axon formation, and branching, while nocodazole treatments result in diminished neurite extension and increased retraction (He et al., 2002; Witte et al., 2008). The altered expression of, or mutations in, microtubule-associated proteins also alter neuron morphogenesis (reviewed in Lasser et al., 2018). For example, the microtubule-associated protein SSNA1 also promotes axonal branching by mediating microtubule nucleation and branching, further highlighting the role of microtubule dynamics play in neuron morphogenesis (Basnet et al., 2018). Neurons must tightly control both microtubule turnover and nucleation to control neurite branching (Basnet et al., 2018; Witte et al., 2008). Both taxol-treatment and SSNA1 overexpression result in increased neurite branching, yet each drive this phenotype via separate mechanisms; Taxol suppresses microtubule turnover, whereas overexpression of SSNA1 induces microtubule nucleation and branching. We find that DIV2 neurons ectopically expressing *TUBA1A*-V409I or -V409A have increased levels of microtubule acetylation, but not increased amounts of microtubule polymer (Figures 1B and 3C). Therefore, our data suggest that V409I/A mutants are unique in that they behave similarly to taxol-treated cells and do not promote increased nucleation compared to WT, but instead result in longer lived microtubules that promote neurite branching.

Our results from modeling V409 mutants in budding yeast reveal the mechanistic origins of the highly stable microtubules in neurons. We find that, compared to WT, *tub1*/*tub3*-V410I/A microtubules have faster polymerization and depolymerization rates, decreased catastrophe frequencies, and increased dynamicity (Figure 4). Dynamicity is calculated as the total change in microtubule length (taking into account states of growth and shrinkage) divided by the change in time (Jordan et al., 1993). It is interesting that despite the *tub1*/*tub3*-V410I/A microtubules undergoing catastrophe less often than WT, they have increased dynamicity values. This suggests that the increase in dynamicity is driven by increased polymerization and depolymerization rates, as opposed to frequent transitions between the states of polymerizing and depolymerizing. Accordingly, V410I/A microtubules reach longer lengths and are longer lived. Assuming these dynamic parameters hold in neurons, longer microtubules that catastrophe less often would result in increased microtubule acetylation, which is what we observe in neurons expressing *TUBA1A*-V409I/A. Our data support a model in which the V409I/A mutants promote excessive neurite branching similar to taxol-treated neurons, but alter microtubule dynamics in a way that is distinct from the mechanisms of taxol.

Microtubule dynamics in cells are reliant on both extrinsic factors that interact with microtubules, as well as intrinsic factors conferred by the tubulin heterodimer (Bodakuntla et al., 2019; Borys et al., 2020; Goodson and Jonasson, 2018; Manka and Moores, 2018; Mitchison and Kirschner, 1984). One way in which tubulin heterodimers intrinsically control microtubule dynamics is by adopting a series of conformations as microtubules polymerize and depolymerize (Chrétien et al., 1995; Mandelkow et al., 1991). Free heterodimer exists in a kinked conformation, which straightens out as the heterodimer is assembled into the microtubule lattice (Buey et al., 2006; Jánosi et al., 1998; Nawrotek et al., 2011; Nogales and Wang, 2006; Ravelli et al., 2004; Rice et al., 2008). We find that V410I/A tubulins have decreased affinity for Stu2 TOG1/TOG2 domains in our *in vitro* binding assays (Figure 5C). Because Stu2 TOG domains preferentially bind to the kinked conformation of the tubulin heterodimer, this raises the possibility that V410I/A mutants push the heterodimer into a straightened conformation. Our finding that V410A increases the rate of tubulin polymerization in the absence of extrinsic MAPs is also consistent with an effect on tubulin’s intrinsic conformation (Figure 6C). We propose a model in which tubulin conformation acts more as a dial than an on and off switch (Figure 7). In this model, limiting the range of conformations a heterodimer adopts is likely to affect the severity of the impact it has on microtubule dynamics. Based off the results of our *in vitro* binding assays between tubulin and Stu2 TOG1/2, V410I appears to have a milder effect on tubulin conformation as compared to V410A (Figure 5C). We interpret this as that V410I can no longer adopt the full range of conformations, but can adopt more of a kinked state than V410A, and thus has less drastic effects on microtubule dynamics. In accordance with our prediction that V410I/A mutants are in a straighter conformation than WT, we find that *tub1*/*tub3*-V410I microtubules, and more significantly -V410A microtubules, have less Stu2 at their plus ends (Figure 5B). Additionally, when Stu2 is conditionally depleted in cells, *tub1*/*tub3*-V410I/A microtubules are maintained while WT microtubules are lost. Together, our data indicate that a consequence of limiting tubulin conformational states is diminished regulation that is typically conferred by Stu2.

**Figure 7.**
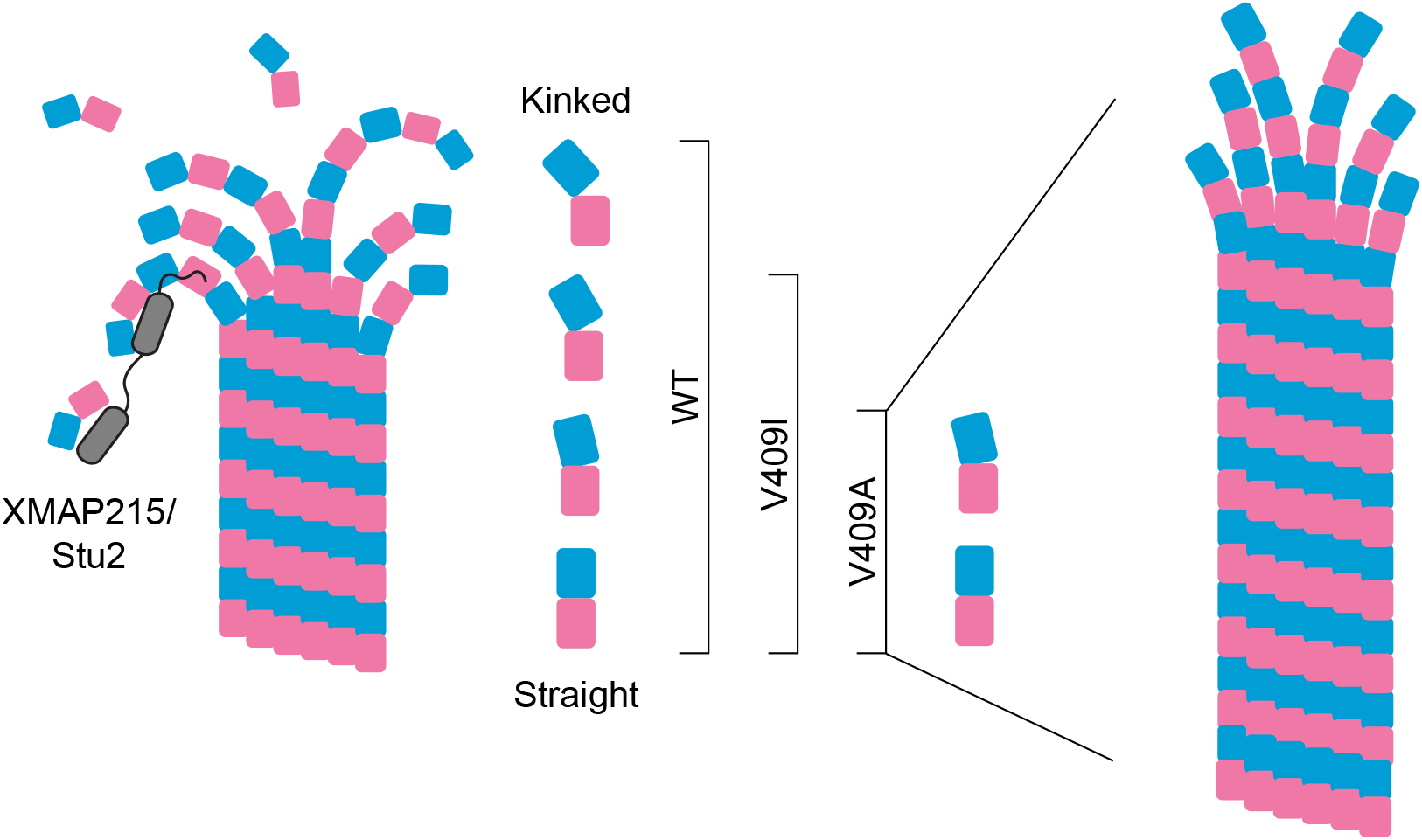
Proposed model for α-tubulin V409I and V409A on tubulin conformations and microtubule assembly. WT tubulin undergoes a series of conformational states as it transitions from a curved free heterodimer into the straight microtubule lattice. Our data support a model in which V409I, and more severely V409A, limit the range of conformations the heterodimer can adopt and push the heterodimer into a straightened state. Additionally, these mutants result in longer-lived microtubules that polymerize faster. We propose that this is due in part to the straighter mutant tubulin being more compatible with assembling into the microtubule lattice, as well as the mutants subverting normal XMAP215/Stu2 regulation.

Tight control over microtubule dynamics is essential for cells, and particularly in neuron morphogenesis and migration as the cytoskeletal must nimbly respond to cues throughout the migratory process (Kapitein and Hoogenraad, 2015). XMAP215/Stu2 works in cells as an extrinsic regulator to finely tune microtubule dynamics (Brouhard et al., 2008; Hahn et al., 2021; Lee et al., 2001; van der Vaart et al., 2011; Vasquez et al., 1994; Zanic et al., 2013). The role of XMAP215 in neurons has been explored primarily in the axon and growth cone. XMAP215’s polymerase activity is required for axon outgrowth and microtubule bundling, and in the growth cone XMAP215 plays a role in linking microtubules and F-actin (Hahn et al., 2021; Lowery et al., 2013; Slater et al., 2019). However, further studies should focus on the role XMAP215 plays in neuron morphogenesis and migration, including in establishing neuron polarity and branching. Beyond XMAP215/Stu2, which preferentially binds to the kinked conformation of tubulin, there are numerous other microtubule-associated proteins that bind to a range of particular tubulin conformations (Brouhard and Rice, 2018). Therefore, we predict that the activity of multiple extrinsic regulators would be impacted by mutations that alter the conformational state of the tubulin heterodimer. How these regulatory mechanisms are impacted by tubulin conformational states and the subsequent effects of these changes on microtubule networks in cells will be a major focus of future studies.

Our results, particularly for the V410A mutant, highlight how tubulin mutants can create unexpected dissonance between extrinsic and intrinsic factors. Compared to WT tubulin, V410A mutant tubulin promotes faster polymerization rates in cells and *in vitro*. It is interesting that in our *in vitro* system, we were unable to identify a concentration at which homogeneous V410A polymerizes from the GMPCPP-stabilized porcine tubulin seeds. However, blending V410A with WT tubulin in a one-to-one ratio supports the assembly of microtubules with significantly increased polymerization rates as compared to WT at the same concentrations of total tubulin. This suggests that V410A may be unable to polymerize from GMPCPP seeds on its own, but it drives fast polymerization when mixed with WT tubulin. Based on our data acquired from neuron cultures, and particularly from budding yeast in which all the α-tubulin in the cell is V410A, we know that this mutant assembles into microtubules that polymerize faster and are longer lived in cells. Why then is V410A able to assemble into microtubules when it is the only source of α-tubulin in the cell, but not when it is the only source of α-tubulin *in vitro*? One model we have considered is that V410A disrupts the delicate equilibrium that exists between free tubulin heterodimer, templated nucleation, and spontaneous nucleation. In cells there are factors such as the γ-tubulin ring complex and various MAPs that promote tubulin nucleation into microtubules. *In vitro*, our yeast tubulin must assemble on its own from GMPCPP-stabilized porcine tubulin seeds. Our *in vitro* dynamics data indicate that when V410A is in a one-to-one blend with WT, the x-intercept is similar to pure WT (Figure 6C). This indicates that the blend would not be expected to nucleate at lower concentrations than WT alone. However, pure V410A may have a lower critical concentration than WT and may shift the equilibrium toward spontaneous nucleation away from stabilized seeds. However, when the mutant is blended with WT, the WT may shift that equilibrium back and promote templated microtubule assembly.

Beyond V410A, interactions between different tubulin heterodimers (such as different mutants or isotypes) is an important question to address both in regards to tubulinopathy disease as well as normal tubulin biology. In tubulinopathy patients that harbor heterozygous mutations, the mutant tubulin comprises at most 50% of either the α- or β-tubulin pool. This underscores the necessity of future work to 1) determine the relative protein constituency of different tubulin isotypes in different cell types, and 2) use blending experiments *in vitro* to recapitulate these relative proportions. Additionally, this is an important consideration for furthering our understanding of how normal tubulin biology works in cells. Using mutants that affect particular intrinsic properties of tubulin (e.g. conformation, GTPase activity, etc.) in various blends with WT tubulin will provide insight into how these different properties work in trans to alter microtubule dynamics, and whether some mutants or isotypes might elicit outsized effects.

We propose a model in which the V409I/A mutants alter the conformational transitions of the heterodimer and push it into a straighter state that drives faster microtubule dynamics and ultimately perturb neuron morphologies and migration (Figure 7). An isoleucine or alanine substitution at the valine 409 residue of *TUBA1A* seems relatively insignificant. However, these subtle changes lead to drastic consequences that scale from altered microtubule dynamics to an underdeveloped neocortex. Strikingly, *TUBA1A*-V409I presents an intermediate phenotype between WT and *TUBA1A*-V409A in each assay we tested, tracking with the observed brain malformations observed in these patients. To date, there are numerous missense, heterozygous mutations in human tubulin genes that are associated with brain malformations, yet the molecular mechanisms driving these tissue level defects remain unknown. Based off our work and that of others, there does not appear to be a consistent mechanism for how *TUBA1A* tubulinopathy mutations affect microtubule networks and neuron cell biology (Aiken et al., 2019; Belvindrah et al., 2017; Gartz Hanson et al., 2016; Keays et al., 2007). The well-established conformational dynamics of tubulin raise the possibility that mutations altering one region of the heterodimer could allosterically affect distant regions, and thus disrupt the normal conformational transitions that heterodimers undergo during assembly and disassembly. Therefore, it is crucial to use quality genetic models to further our understanding of the molecular mechanism of these patient-associated mutations. Additionally, our work highlights the need for more detailed studies into how different regions of the tubulin heterodimer impact tubulin conformational states, and how this affects extrinsic regulation that ultimately result in altered microtubule dynamics. Investigating how subtle alterations in specific regions of the heterodimer can have long ranging effects across microtubule networks will be a primary focus of future work.

## ACKNOWLEDGEMENTS

We are grateful to members of the Moore and Dr. Emily Bates (CU AMC) labs for helpful discussions. This work was supported by T32 GM136444 (K.J.H.); Center for Neuroscience Pilot Award from the Colorado Clinical and Translational Sciences Institute (funded by NIH UL1TR002535), Boettcher Webb-Waring Biomedical Research Program Award from the Boettcher Foundation, and the Children’s Hospital Colorado Program in Pediatric Stem Cell Biology (S.J.F.); and NIH R35GM136253 (J.K.M.).

## MATERIALS AND METHODS

### Animals

Animal research was performed following regulations approved by the Institutional Animal Care and Use Committee at the University of Colorado School of Medicine. All mouse tissue experiments described below were from *C57BL6/J* wild-type pregnant mice (Jackson Laboratories Strain #000664). All dissociated neuron cultures described below were prepared from Sprague-Dawley rats. Dissociated cultures were obtained from both male and female pups.

### TUBA1A plasmid vectors

Plasmids used in this study are listed in Supplementary Table 1. The plasmids were constructed using the methods described in Aiken et al. 2018. Briefly, the coding region of the human *TUBA1A* gene was cloned into the multiple cloning site of the pCIG2 plasmid that express cytoplasmic GFP from an internal ribosome entry site (IRES) (Hand et al., 2005). QuikChange Lightning Site-Directed Mutagenesis Kit (Agilent Technologies, Santa Clara, CA) was used to introduce either the p.V409A or p.V409I substitution into TUBA1A. All plasmids have the 6X-His tag inserted between residues I42 and G43. Mutants were confirmed by sequencing. Oligos used for plasmid constructions are listed in Supplementary Table 3.

### Primary Cortical Neuron Cultures

The frontal cortex was dissected from postnatal days 0-2 male and female neonatal Sprague Dawley rats. Cortices were digested at room temperature for 1 hour using papain solution composed of 20 units/ml papain (Worthington Biochem, LS003126), 1.5 mM CaCl_2_ (Sigma Aldrich, 223506), 0.5 mM EDTA (Fisher Scientific, BP118-500), 1 mM NaOH (Fisher Scientific, BP359-500), and 0.2mg/ml cysteine (Sigma Aldrich, C6852) in 10 ml saline. Digested cortices were washed six time with Minimal Essential Media (MEM) (Life Technologies, Carlsbad, CA, 11090-081). Cortices were dissociated with a wide-bore fire-polished pipette, followed by dissociation with a narrow-bore polished pipette. pCIG2-TUBA1A-His6 (either EV, WT, or mutant) plasmids (4µg) were introduced to 4×10^6^ dissociated neurons using Amaxa rat nucleofector kit (Lonza Bioscience, VPG-1003). Amaxa-nucleofected neurons plated at 500,000 cells per 35mm poly-D-lysine-coated glass bottom dish (WillCo Wells, HBST-3522) in MEM containing 10% Fetal Bovine Serum (FBS) (Sigma Aldrich, F4135), 1% penicillin/streptomycin (Gibco, 15070-063), and 25 µM L-glutamine (Gibco, 25030081). Following 2 hours in culture, media was replaced with supplemented FBS. At 24 hours after plating, media was replaced with Neurobasal-A (NBA) (Life Technologies, 10888-022), 2% B27 (Gibco, 17504-044), and 10 µM uridine/fluoro-deoxyuridine (Sigma Aldrich, U3003/F0503). Neurons were maintained in a 37°C humidified incubator with 5% CO_2_.

### Neuron Immunocytochemistry

Days in vitro (DIV) 2 or DIV3 primary cortical neurons were washed with phosphate buffered saline (PBS) (Gibco, 10010023) followed by PHEM buffer, which contains 60 mM PIPES (Sigma Aldrich, P6757), 25 mM HEPES (Sigma Aldrich, H3375), 10 mM EGTA (Sigma-Aldrich, E3889), and 2 mM MgCl_2_ (Acros, 223211000). Soluble tubulin dimers were extracted using 0.1% triton-X (Sigma Aldrich, T9284) with 10 µM taxol (LC Labs, P-9600) and 0.1% DMSO (Fisher, M-1739) in PHEM buffer. Cells were fixed with 4% paraformaldehyde (PFA) (Sigma Aldrich, 158127) and 0.1% glutaraldehyde (Sigma Aldrich, G7776) in PBS for 10 minutes at room temperature, washed with PBS, then blocked with blocking buffer, 3% BSA (Fisher, 50-253-893) and 0.2% Triton-X in PBS. Cells were reduced with 10 mg/ml sodium borohydride (Fisher, S678-10) in an equal mixture of PBS and methanol (Acros, 177150050) for 7 minutes at room temperature, then washed three times with PBS. Cells were incubated with blocking buffer for 20 minutes at room temperature. Immunostaining was performed using a primary antibody directed against 6X-Histidine (Invitrogen, 4A12E4 37-2900; 1:500), beta-II tubulin (Abcam, ab179512; 1:500), acetylated tubulin (Sigma, T7451; 1:1000), tyrosinated tubulin (Sigma, MAB1864-I; 1:1000). Primary antibody was diluted in blocking buffer and incubated overnight at 4°C in a humidified chamber. After primary antibody staining, cells were washed three times with PBS. The secondary antibodies goat anti-rabbit IgG Alexa Fluor 568 (Thermo, A-11011), goat anti-rat IgG Alexa Fluor 568 (Thermo, A-11077), or goat anti-mouse IgG Alexa Fluor 647 (Thermo, A32728) were diluted 1:200 in blocking buffer and 1% normal goat serum (Vector Laboratories, S-1000-20) and incubated for 2 hours at room temperature in a dark container. Cells were sealed with glass coverslips and aqueous mounting media containing DAPI (Vector Laboratories, H-1200), then imaged on a spinning disk confocal microscope with a 40x oil objective (see details below). Statistical analysis between multiple groups were analyzed by two-way ANOVA and analyzed *post hoc* by Tukey test.

### In utero electroporation

*In utero* electroporations were performed on E14.5 embryos of *C57BL6/J* wild-type pregnant mice. Plasmid DNA was endotoxin-free and prepared with 0.1% fast green dye and had a final concentration of 1µg/µl in TE buffer (10mM Tris base and 1mM EDTA, pH 8.0). Plasmid DNA was injected into the lateral ventricles of the exposed embryos and electroporated with 5 pulses at 50V for 100ms pulses each, separated by 950ms. All embryos developed to E18.5, then brains were dissected and fixed overnight at 4°C in 4% paraformaldehyde. A vibrating microtome (VT1200S; Leica, Buffalo Grove, IL) was used for 50µm thick coronal sections. Sections were mounted on glass slides, then imaged on a spinning disk confocal microscope with a 20x objective. Analysis includes at least three animals, both male and female, per condition.

### Neuron cold temperature shock

The media of DIV2 neuron cultures (as described above) was replaced with Neurobasal-A (NBA) (Life Technologies, 10888-022), 2% B27 (Gibco, 17504-044), and 10 µM uridine/fluoro-deoxyuridine (Sigma-Aldrich, U3003/F0503). The dishes were then placed at 4°C for the desired time. At each time point, dishes were removed from 4°C and placed on ice while following the fixing protocol described above.

### Neuron morphology and post-translational modifications analysis

All images were analyzed using ImageJ (National Institute of Health) and the NeuronJ plugin (Meijering et al., 2004). Neuron processes were defined as primary, secondary, or tertiary, then quantified for each cell analyzed. Post-translational modification fluorescent signals were quantified from each cell by measuring line scans along processes. For consistency within each individual cell, the line scans were first traced using cytoplasmic GFP signal, saved, and then used to measure signal from the immunofluorescent staining for the tubulin PTMs in different channels. A two-way ANOVA analyzed *post hoc* by Tukey test was used for statistical analyses between multiple groups.

### Yeast microtubule dynamics

Microtubule dynamics were analyzed in log-phase, pre-anaphase yeast cells in which all the α-tubulin in the cell was either WT, V410I, or V410A. The mutants were integrated at the native *TUB1* and *TUB3* loci. Microtubule plus-ends were identified by Bik1-3GFP. Images were collected on a Nikon Ti-E microscope equipped with a 1.45 NA 100× CFI Plan Apo objective, piezo electric stage (Physik Instrumente, Auburn, MA), spinning disk confocal scanner unit (CSU10; Yokogawa), 488-nm and 561-nm lasers (Agilent Technologies, Santa Clara, CA), and an EMCCD camera (iXon Ultra 897; Andor Technology, Belfast, UK) using NIS Elements software (Nikon). Cells were grown asynchronously to early log phase in nonfluorescent media and adhered to slide chambers coated with concanavalin A. Slide chambers were sealed with VALAP (Vaseline, lanolin and paraffin at 1:1:1). Z-series consisting of a 7µm range at 0.4µm steps were acquired every 4 seconds for 10 minutes at 30°C. Astral microtubule lengths were measured in each frame as the distance from the spindle pole body to the microtubule plus-end. Cell genotypes were blinded for analysis. Statistical analysis was done using a one-way ANOVA followed by a Tukey test to correct for multiple comparison tests.

### Stu2 localization

Analysis was done in log-phase, pre-anaphase budding yeast cells expressing either *tub1/tub3*-WT, V410I, or V410A, as well as Stu2-3GFP. The tubulin mutations and the fluorescent tagging of Stu2 were done at the endogenous loci. Short time-lapse images, every 5 seconds for 30 seconds, were collected on a spinning disc confocal microscope with a 100x oil objective, using Z-series consisting of 0.4µm steps over a total range of 7µm. Stu2-3GFP fluorescence intensity was measured at the plus-ends of growing astral microtubules by defining a 182nm^2^ region. A region of the same size and adjacent to the Stu2-3GFP foci was also measured and subtracted from the Stu2-3GFP intensity value to account for background fluorescence. Statistics represent values from a one-way ANOVA corrected for multiple comparison by a Tukey test.

### GST-TOG1/2 purification

Purification of GST-TOG1/2 followed previously described methods and is described briefly below (Reusch et al., 2020; Widlund et al., 2012). The pGEX-6P-1 Stu2 1-590 plasmid was transformed into BL21 cells and colonies were grown on an LB plate containing 100µg/ml carbenicillin and 15µg/ml chloramphenicol at 37°C. A colony was inoculated overnight at 37°C in MDAG-135 media (25mM Na_2_HPO_4_, 25mM KH_2_PO_4_, 50mM NH_4_Cl, 5mM Na_2_SO_4_, 2mM MgSO_4_, 0.2x metals, 0.35% glucose, 0.1% aspartate, 200µg/ml each of 18aa (no C, Y)) containing 100µg/ml carbenicillin and 15µg/ml chloramphenicol (Studier, 2005). The following day the colony was diluted 500-fold in 1L of Terrific Broth media (Fisher, BP2468-2) with 100µg/ml carbenicillin and 15µg/ml chloramphenicol and grown at 37°C until the OD_600_ reached approximately 0.5. The cultures were shaken at 18°C for 1 hour, then induced with 0.2mM isopropyl β-_D_-1-thiogalactopyranoside for 18 hours (Fisher, BP1620-1). Following induction cells were pelleted at 6,200xg for 10 minutes at 6°C and resuspended in a 1:1 ratio with a buffer consisting of 2X phosphate-buffered saline (PBS), 1mM dithiothreitol (DTT), 20µl benzonase (25U/µl), and 1X protease inhibitors (2 cOmplete EDTA-free Tablets per 100ml; Roche, 04693132001). The cells were lysed by two passes through a microfluidizer at 1,500 bar. Cell lysate was then clarified by spinning at 12,000rpm for 30 minutes at 4°C.

A 5mL GSTrap HP column (GE, 17-5282-01) was preequilibrated with 5 column volumes (CV) of wash buffer (2X PBS, 1mM DTT) at 1mL/min. The clarified lysate was then loaded onto the column at 0.5mL/min. The following washes were all done at 1mL/min. The column was washed with 10CV of wash buffer with 0.1% Tween 20, then 2CV of 2X PBS with 5mM ATP and 10mM MgCl_2_. The column incubated in this buffer for 20 minutes. The column was then washed with 5CV of 6X PBS, followed by 5CV of wash buffer. GST-TOG1/2 was eluted from the column in 1mL fractions into a deep 96-well plate with wash buffer plus 5mM reduced glutathione, pH 8.0. The eluted fractions were run on a gel and stained with Coomassie. Fractions containing GST-TOG1/2 were pooled and dialyzed against three changes (after 2h, overnight, then another 2h) of coupling buffer (100mM NaHCO3, 100mM NaCl, pH 8.2) at 4°C. The dialyzed sample was concentrated in 0.5ml centrifuge concentrating filters (Sigma Z677108) that had been preequilibrated with coupling buffer. GST-TOG1/2 was concentrated to 14.5µM and stored in 25µl aliquots at -80°C.

### Yeast tubulin purification

Purification of yeast tubulin was based on previously described methods with slight modifications (Minoura et al., 2013). Protease deficient yeast cells (JEL1) were transformed with pRS426-GAL-Tub1-internal 6X-His and pRS424-GAL-Tub2 (untagged). tub1-V410I and -V410A mutants were constructed in the pRS426-GAL-Tub1-internal 6X-His plasmid with QuikChange XL. Cells were grown in five or six 5ml cultures of selection media (-ura -trp) supplemented with 2% glucose for either three or two days, respectively, at 30°C. 5ml cultures were then transferred to 50ml cultures of the same media to grow for 24 hours. Cultures were then transferred to 1L cultures of YPGL, consisting of 10g yeast extract (Fisher, BP1422), 20g peptone (Fisher, BP1420), 30ml glycerol (Fisher, G33), and 33ml lactate (Sigma, L1375). Yeast extract, peptone, and 850ml ddH_2_O was combined and autoclaved the day before it was needed. The glycerol and lactate were added immediately after media came out of the autoclave, then allowed to cool at room temperature overnight. After 20-24 hours of growing in 1L cultures and the OD600 was between 5.0-9.0, 1L of cells was induced with 20g galactose (Chem-Impex, 01449) for five hours. Cells were spun down at 4°C for 15 minutes at 4000 RPM (3040 RCF) in a J6 centrifuge (Beckman Coulter Life Sciences). Supernatant was discarded and pellets were stored in 15ml conical tubes in -80°C. This process was done three times for a total of approximately 100g of cells before moving on to purification.

Cell pellets were thawed on ice and combined with cold lysis buffer (50mM HEPES, 500mM NaCl, 10mM MgSO_4_, 30mM Imidazole) plus freshly added 50µm GTP (Sigma G8877) and 1mM PMSF (Acros Organics, 215740050) to a total of approximately 150ml. While cells were thawing, the microfluidizer (Microfluidics, M-110P) was packed with ice to cool. Cells were passed through the microfluidizer four times at 27,000 PSI with 10 minutes on ice between each pass. The cells were chased with 50ml lysis buffer supplemented with 50µM GTP and 1mM PMSF. The lysed cells were clarified in an Avanti J-26S XPI centrifuge (Beckman Coulter Life Sciences) at 41,000 RCF for 30 minutes at 4°C. The pellets were discarded and the supernatant was applied to a 5ml His60 Ni column (Takara, 635680) that was pre-equilibrated with 20 column volumes of lysis buffer. The column was washed with 10 column volumes of lysis buffer supplemented with 50µM GTP, then with 10 column volumes of wash buffer (25mM PIPES pH 6.9, 1mM MgSO_4_, 30mM Imidazole, 200mM NaCl) with 50µM GTP. Protein was eluted from the column with 10 column volumes of elution buffer (25mM PIPES pH 6.9, 1mM MgSO_4_, 300mM Imidazole, 200mM NaCl) with 50µM GTP into 2 x 5ml samples followed by 4 x 10ml samples. Samples from the wash and elution steps were run on a gel and Coomassie staining was used to determine which fractions contained tubulin. Elution fractions with tubulin were pooled and 20% glycerol was added before flash freezing the samples in liquid nitrogen and storing at - 80°C.

Elution fractions from the Ni column were thawed in a room temperature water bath and 10µl of nuclease (Pierce, 88701) was added per 25ml of sample, then sat for one hour at room temperature. Two HiTrap Q HP 1ml columns (GE, 17-1153-01) were strung together on an AKTA FPLC (GE) and equilibrated with 10 column volumes (20ml) of ddH_2_O, 5 column volumes of Buffer A (25mM PIPES pH 6.9, 2mM MgSO_4_, 1mM EGTA), 5 column volumes of Buffer B (25mM PIPES pH 6.9, 2mM MgSO_4_, 2mM EGTA, 1M NaCl), and 10 column volumes of Buffer A, all run at 1ml/min. The column was then equilibrated at 2ml/min with 12.5 column volumes of 80% Buffer A and 20% Buffer B, both supplemented with 50µM GTP. Sample from Ni column was injected into a 50ml Superloop that was washed with diH_2_O followed by Buffer A. The sample was applied to the column at 0.5ml/min using Buffer A. Once the sample was loaded the tubulin was eluted with 30 column volumes of Buffer A and a 20-60% gradient of Buffer B, both supplemented with 50µM GTP. Elution was collected in 0.5ml fractions in a 96 deep-well plate. Elution fractions were run on a gel and Coomassie staining determined which fractions had pure tubulin and would be used to concentrate the protein.

Amicon ultra 0.5ml centrifuge concentrating filters with a 10kDa cutoff (Sigma, Z677108) were prepared with two rounds of 0.4ml 1% Triton X-100, ddH_2_O, and PEM (0.1M PIPES, 1mM EGTA, 1mM MgCl_2_) spinning at 800 RCF for 5 minutes at 4°C. Up to 0.450ml of pure tubulin sample was applied to the concentrating filters and spun at 4°C at 800 RCF for anywhere between 2-5 minutes, depending on the concentration of the sample. The concentration was checked after each spin on a Take3 Micro-volume Plate (BioTek) using 260nm and 280nm absorbance to ensure that the concentration was increasing but aggregates were not forming. Samples were constantly kept on ice or at 4°C and concentrated to 2.2µM (tub1-V410I), 2.3µM (tub1-V410A), and 3µM (TUB1). Concentrated protein samples were run through a Zeba spin 2ml desalting column (Thermo, 89890) that was prepared by spinning at 1000 RCF for 2 minutes at 4°C to remove storage buffer, then washed three times with 0.4ml PEM. Up to 0.7ml of concentrated sample was added to each desalting column, then spun down at 1000 RCF for 2 minutes at 4°C. The flow-through sample was supplemented with 50µM GTP, aliquoted into 20µl samples that were flash frozen in liquid nitrogen, and stored at -80°C. At least two separate rounds of purifications were used as technical replicates for *in vitro* assays that used this purified tubulin.

### TOG binding assay

Pierce Glutathione Magnetic Agarose Beads (Thermo, 78602) were used to separate purified GST-TOG1/2 (above) and anything bound in complex with TOG1/2 from the rest of the reaction. 100µl of bead slurry (equates to 25µl settled beads) were equilibrated twice with 40µl wash buffer (125mM Tris-HCl, 150mM KCl, 1mM DTT, 1mM EDTA, pH 7.4) by vortexing for 10 seconds and removing buffer by placing tube on magnetic stand. In separate tubes, 100nM tubulin (WT, V410I, or V410A; purification described above) was mixed with a range of GST-TOG1/2 concentrations (from 9.8nM to 5µM) at 4°C for 15 minutes. Reaction was added to the equilibrated beads and rotated at 4°C for 1 hour. Tubes were placed on magnetic stand and supernatant was collected. The supernatant was prepared with Laemmli sample buffer and β-mercaptoethanol and loaded onto a 10% SDS-PAGE gel. The gel was transferred to a PVDF membrane (Millipore, IPFL85R), blocked with a PBS Blocking Buffer (LI-COR, 927-70001) at room temperature for 1 hour, and probed with mouse anti-α-tubulin (4A1; 1:100) or mouse anti-β-tubulin (E7; 1:100) overnight at 4°C. The following day the membranes were washed once for 5 minutes with 1X PBS, then incubated at room temperature in the dark for 1 hour with goat anti-mouse-680 (LI-COR, 926-68070; 1:15,000). Membranes were washed twice with 1X PBST, then once with 1X PBS before imaging on an Odyssey Imager (LI-COR, 2471). Both α- and β-tubulin band intensity was analyzed using Image Studio v5.2 (LI-COR). Tubulin band intensity represents the amount of tubulin in solution that is not bound to GST-TOG1/2.

### Stu2 depletion and yeast immunofluorescence

Stu2 depletion strains originally constructed from Kosco et al. were graciously gifted from Dr. Tim Huffaker (Kosco et al., 2001). Strains were modified to mutate the endogenous *TUB1* locus to either V410I or V410A using site-directed mutagenesis. Cells were grown to log-phase, then induced with 500µM cupric sulfate for 1.5 hours at 30°C. The following fix and staining protocol was adapted from (Miller, 2004) as described. Cells were fixed with 3.7% formaldehyde (Sigma-Aldrich, 252549) at 30°C for 2 hours, then spun down at 3000RPM for 3 minutes. Pellet was washed twice with wash buffer (40mM KPO_4_, pH 6.5), then stored overnight at 4°C. The pellet was washed twice with wash buffer plus 1.2M sorbitol. Fixed cells were digested with 10µl of 20T 50mg/ml zymolyase (Nacali Tesque, 07663-91) and 15µl β-mercaptoethanol (Sigma-Aldrich, M3148) for 45 minutes at 37°C. Digested cells were spun down at 2000RPM for 3 minutes, washed once with wash buffer plus 1.2M sorbitol, then resuspended in 20µl wash buffer plus 1.2M sorbitol. 20µl of cells were spotted onto each well of a Teflon coated 10 well slide (Polysciences, 18357) that had been pre-treated with 10ng/µl poly-L-lysine. Cells adhered for 10 minutes at room temperature. Liquid was aspirated off before immediately permeabilizing the cells in a coplin jar of cold methanol for 6 minutes, followed by immersing the slide in cold acetone for 30 seconds. Cells were blocked at room temperature for 1 hour in blocking buffer (1X PBS + 0.5% BSA), then incubated overnight at 4°C in a humid chamber with mouse anti-α-tubulin (4A1; 1:100 in blocking buffer). Wells were washed 4 times for 10 minutes with blocking buffer, then incubated at room temperature for 1 hour in a dark humid chamber with goat anti-mouse IgG-Alexa488 (Invitrogen, A11001). Cells were washed 4 more times with blocking buffer. DAPI mounting solution (Vector Laboratories, H-1200) was added to the slide, then imaged on widefield microscope with a 100x oil objective. Statistical analysis was done using a two-way ANOVA corrected for multiple comparison by a Tukey test.

### Yeast growth assays

For liquid growth assays, saturated cultures were diluted 1:500 in rich media. In a 96-well plate 200µl/well was used. OD600 values were measured on an Epoch 2 Microplate Spectrophotometer (BioTek #EPOCH2NS) every 5 minutes for at least 19 hours at 30°C with orbital shaking. The doubling time was calculated by fitting the measured OD600 values to an exponential curve using a MATLAB code that has previously been described (Fees et al., 2016). The doubling times have been normalized to WT cells. A two-way ANOVA was used to compare across multiple groups and corrected *post hoc* by a Tukey test.

### 4°C temperature shock in yeast

Cells expressing either TUB1/TUB3, tub1/tub3-V410I, or tub1/tub3-V410A along with GFP-Tub1, GFP-Tub1-V410I, or GFP-Tub1-V410A, respectively, were grown overnight at 30°C in rich liquid media. The following day the cells were diluted in fresh media and grown to log phase. The log phase cultures were then transferred to a shaker at 4°C for the indicated amount of time. Cells were fixed at 4°C with 3.7% formaldehyde and 0.1M KPO_4_ for three minutes. Cells were pelleted on a desktop centrifuge and supernatant was discarded. The pellet was resuspended in quencher solution (0.1% Triton-X, 0.1M KPO_4_, 10mM ethanolamine). The cells were pelleted and washed twice with 0.1M KPO_4_. Imaging chamber slides were coated with concanavalin A and washed with 0.1M KPO_4_ before loading fixed cells into the chambers and sealed with VALAP (Vaseline, lanolin, and paraffin at a 1:1:1 ratio) (Fees and Moore, 2018).

### In vitro dynamics analysis using interference reflection microscopy

Interference reflection microscopy was used to measure microtubule dynamics of unlabeled purified tubulin. Chambers were assembled with two different sized coverslips, 22mm x 22mm and 18mm x 18mm, that were silanized and plasma cleaned. Three strips of single ply parafilm were melted between the coverslips to create two chambers on a custom-made stage insert. Anti-rhodamine antibody (Life Technologies, A6397) was diluted 1:100 in cold BRB80 (80mM PIPES, 1mM MgSO_4_, 1mM EGTA), flowed into the chamber, then sat at room temperature for five minutes. The antibody was then washed out with room temperature BRB80. Chambers were flushed with 1% pluornic-F127 in BRB80 and incubated in this buffer for five minutes, then washed out with room temperature BRB80. The stage insert was placed in the scope enclave to equilibrate at 30°C for 30 minutes. GMPCPP-stabilized microtubule seeds that attached to the anti-rhodamine antibody on the coverslips were flowed into the chamber where they settled for 30 seconds before washing out with room temperature BRB80. The chambers were washed with reaction buffer that contained no tubulin (1X PEM, 0.1% methyl cellulose, 1mM GTP, 0.1mg/ml BSA) before adding the reaction buffer along with 0.3µM-0.9µM tubulin. The chamber was then sealed with VALAP. The reaction in the chamber was equilibrated on the widefield microscope for 20 minutes at 30°C to prevent drifting during imaging. Following the equilibration period, images were collected every second for one hour in the interference reflection microscopy channel (MAHAMDEH et al., 2018). The 561nm channel was also collected for the first frame to mark the GMPCPP-stabilized seeds. ImageJ was used for analyzing microtubule dynamics by generating kymographs of individual microtubules.

---

**Supplementary Table 1.** Plasmids.

| Plasmid ID | Plasmid Name | Source |
| --- | --- | --- |
| pJM0157 | pGEX6p1-GST-STU2 1-590 | Widlund et al. 2012 |
| pJM0295 | pCIG2 | Hand et al. 2005 |
| pJM0288 | pRS426-GAL-Tub1-internal 6Xhis | Johnson et al. 2011 |
| pJM0289 | pRS424-GAL-Tub2 (untagged) | Johnson et al. 2011 |
| pJM459 | Stu2-3GFP | This study |
| pJM0469 | TUB1-pRS306 integrating plasmid | Aiken et al. 2020 |
| pJM507 | Tub1-V410I-pRS306 integrating plasmid | This study |
| pJM560 | Tub1-V410A-pRS306 integrating plasmid | This study |
| pJM0546 | pCIG2-TUBA1A-6XHis | Buscaglia et al. 2020a |
| pJM0558 | pRS426-GAL-Tub1(V410A)-internal 6Xhis | This study |
| pJM0559 | pRS426-GAL-Tub1(V410I)-internal 6Xhis | This study |
| pJM0612 | pCIG2-TUBA1A(V409I)-int-6XHis | This study |
| pJM0642 | pCIG2-TUBA1A(V409A)-int-6XHis | This study |

**Supplementary Table 2.**
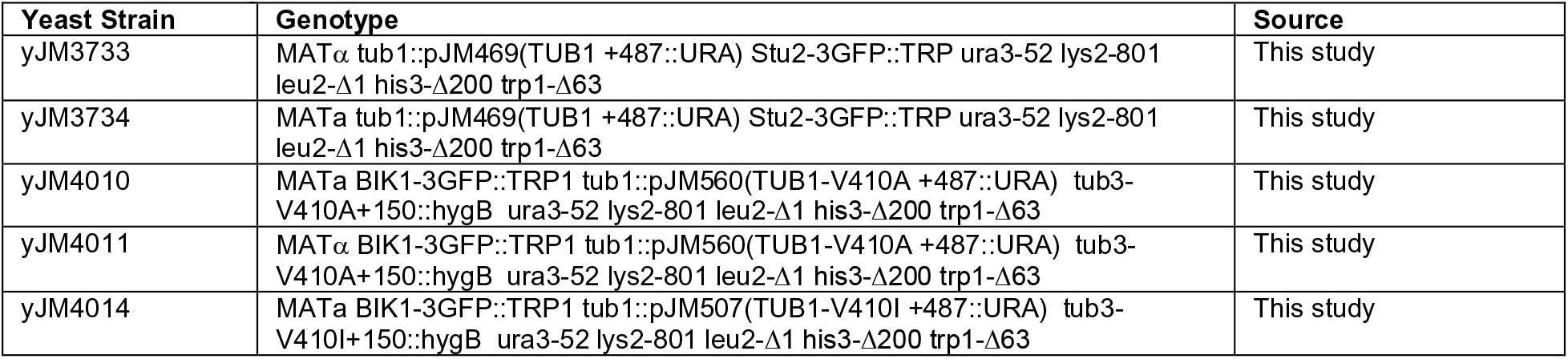
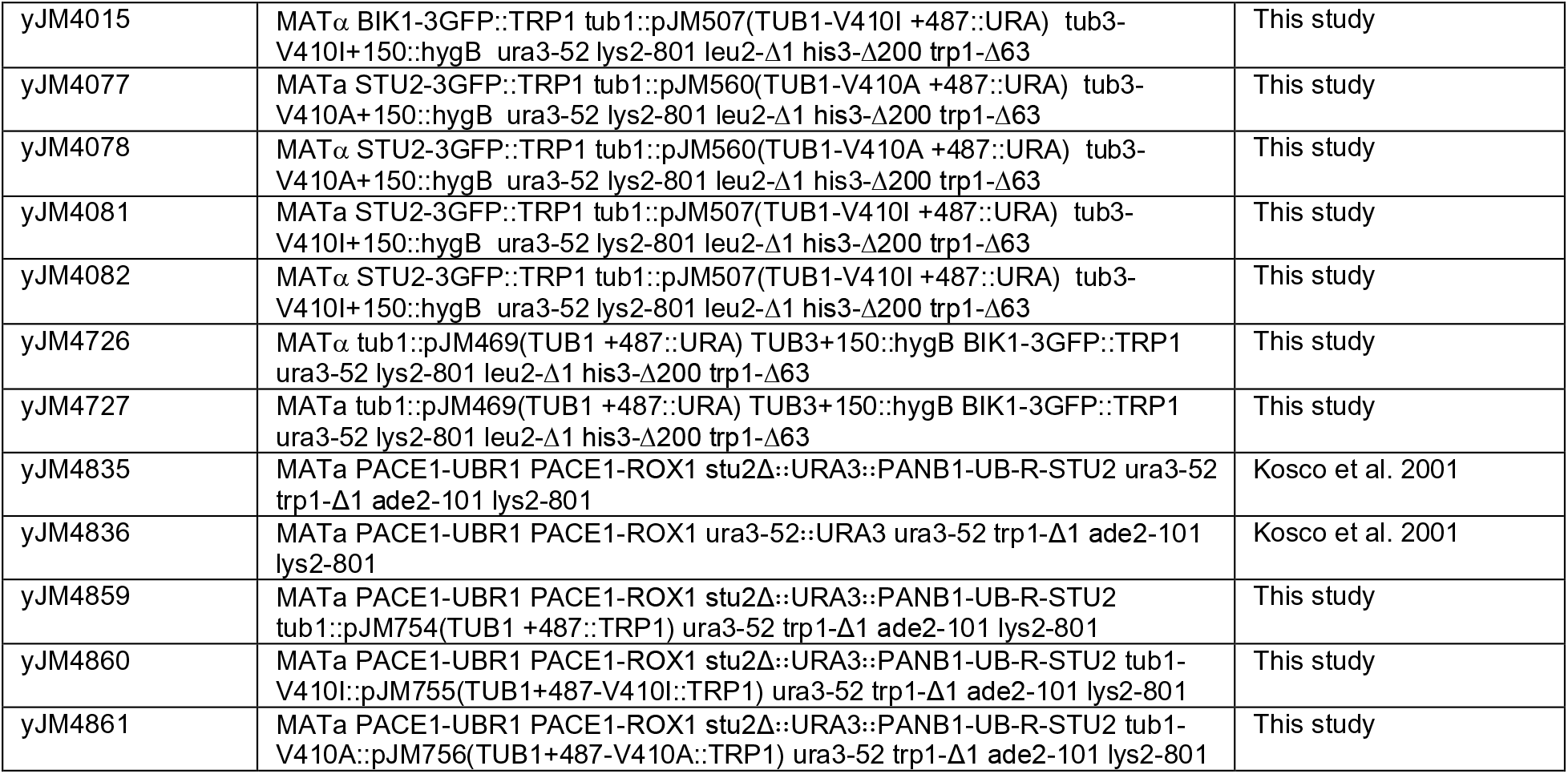
Yeast strains.

**Supplementary Table 3.**
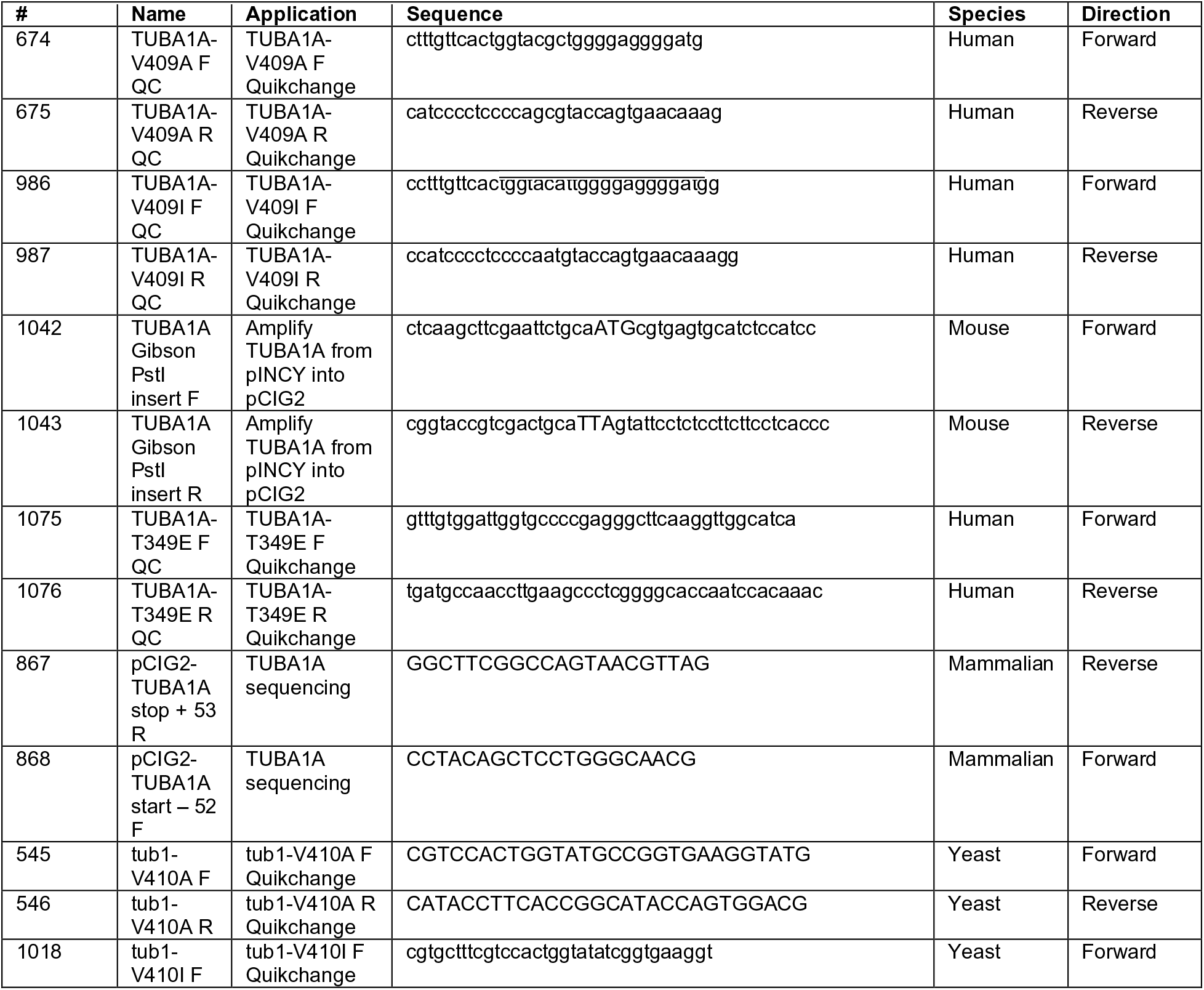
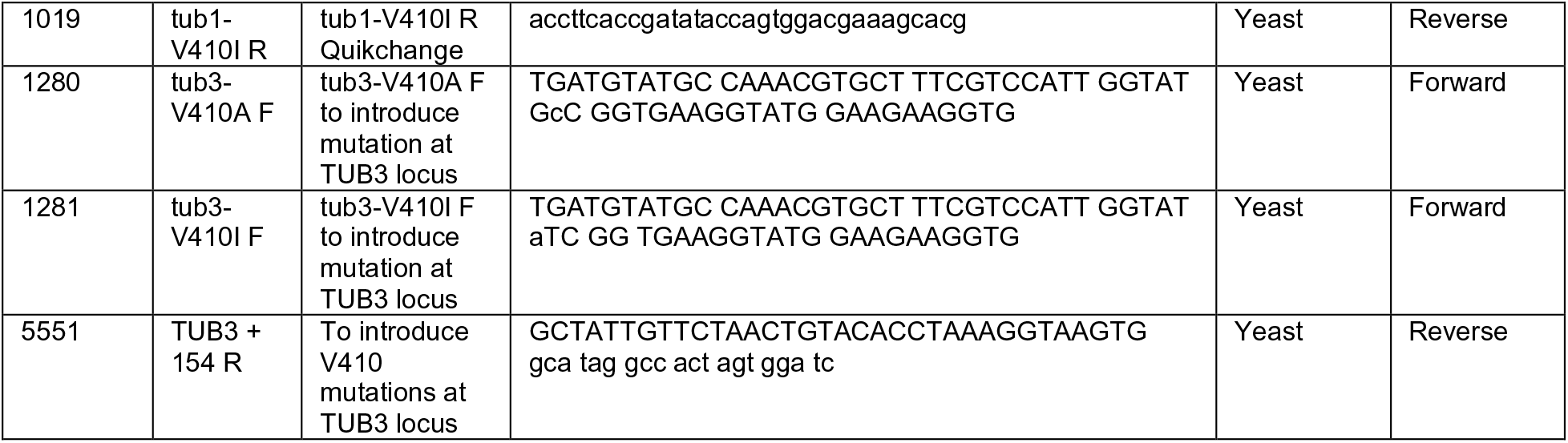
Oligo sequences (5’ to 3’)

**Supplemental Figure 1.**
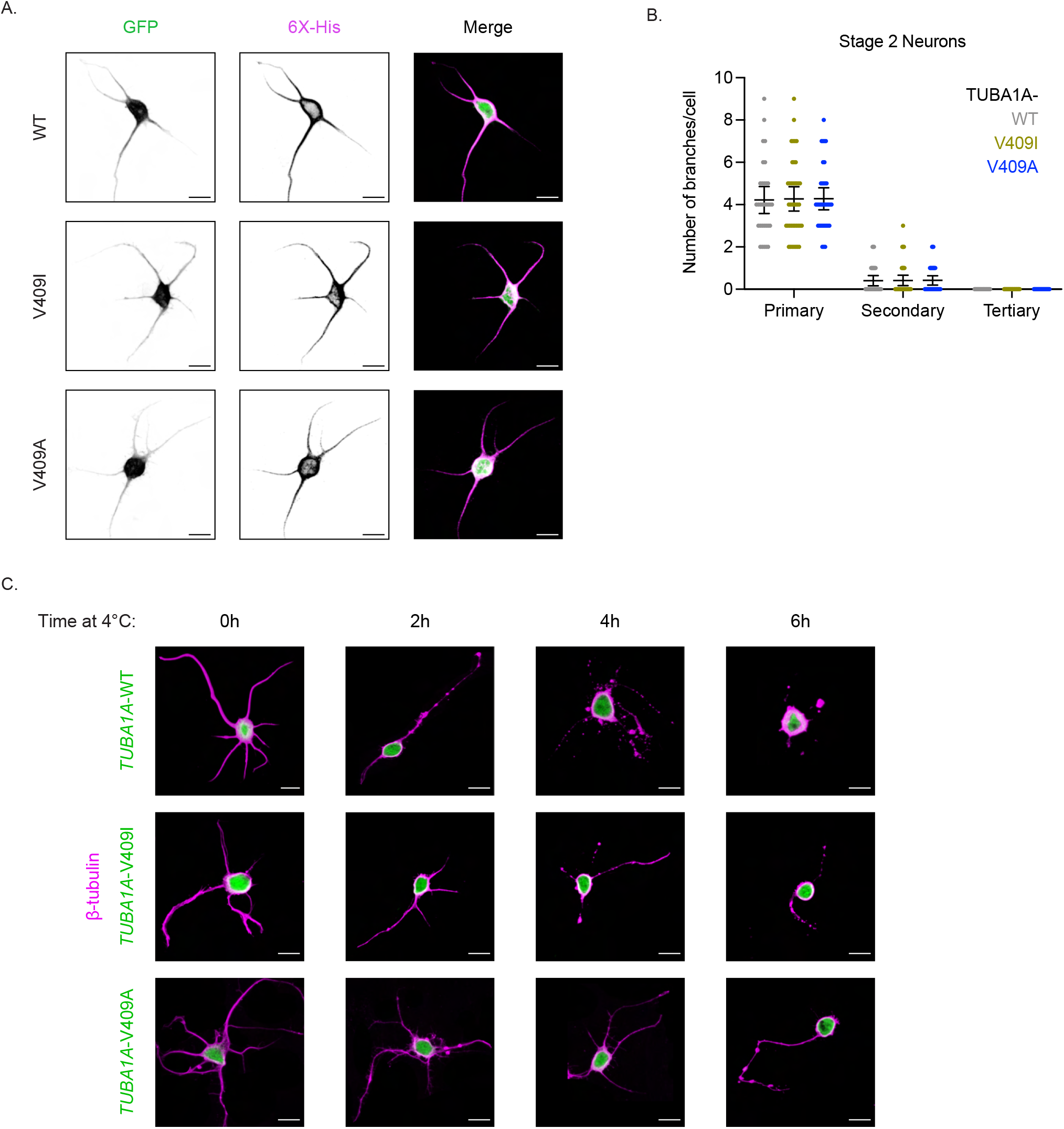
Effect of *TUBA1A*-V409I/A on neuronal morphologies. **A)** Representative DIV1 cortical neurons expressing GFP and 6X-His tagged *TUBA1A*-WT, -V409I, and -V409A. Scale bar = 10μm. At least 23 neurons were imaged for each condition. **B)** Quantification of number of primary, secondary, and tertiary branches in each condition. Each dot represents a single cell. Bars are mean ± 95% confidence interval. **C)** Representative merge images of neurons expressing the above plasmids, exposed to 4°C for indicated time, stained with TUBB2A/B in magenta, and expressing cytoplasmic GFP in green. Scale bar = 10μm.

**Supplemental Figure 2.**
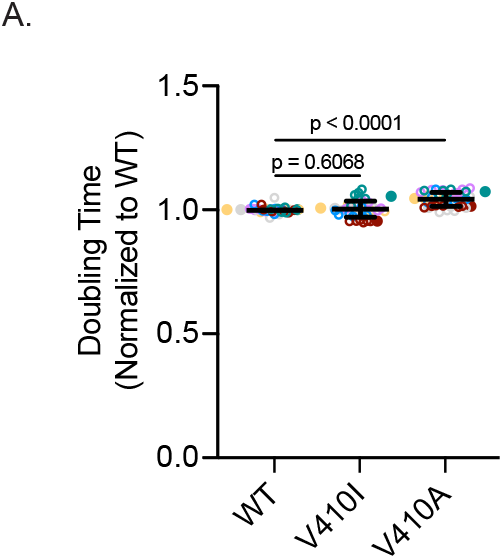
Fitness assay for V410I/A yeast mutants. **A)** Doubling times of cells expressing the indicated mutation at the genomic *TUB1* locus. 6 samples containing 2 different isolates for each strain were tested in 3 independent experiments. Bars are mean ± 95% confidence interval.

**Supplemental Figure 3.**
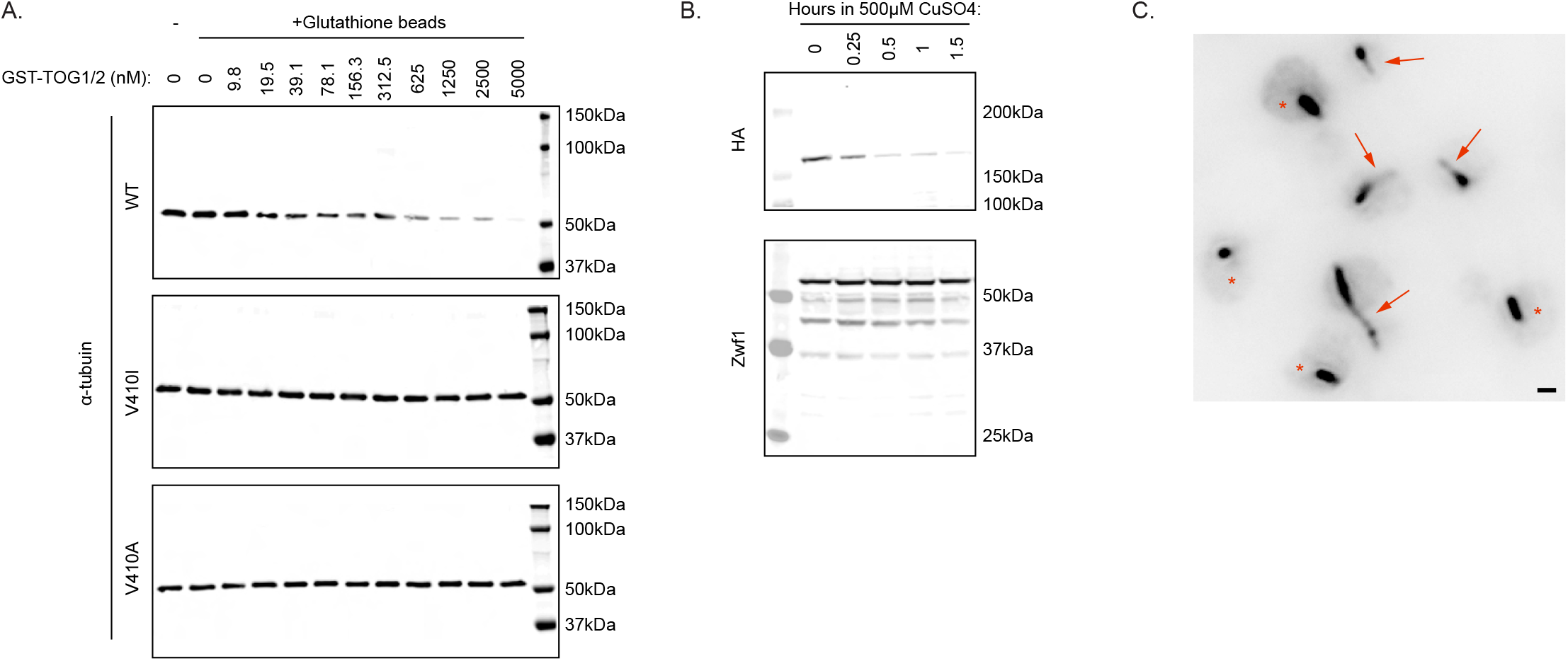
V410I/A mutants affect Stu2 interactions and regulation. **A)** Representative western blots of WT, V410I, and V410A samples probed for α-tubulin in the absence of glutathione beads, as well as in the presence of glutathione beads and increasing concentrations of GST-TOG1/2. **B)** Representative western blot of WT cells induced with 500μM CuSO_4_ for indicated number of hours. Blots were probed with HA and Zwf1 antibodies. **C)** Large field representative image of WT cells in the absence of 500μM CuSO_4_ stained for α-tubulin. Red arrows represent cells with at least one astral microtubule. Red stars represent cells without an astral microtubule. Scale bar = 1μm.

## References

Aiken J, Buscaglia G, Aiken AS, Moore JK, Bates EA. 2020. Tubulin mutations in brain development disorders: Why haploinsufficiency does not explain TUBA1A tubulinopathies. Cytoskeleton 77:40–54. doi:10.1002/cm.21567

Aiken J, Moore JK, Bates EA. 2019. TUBA1A mutations identified in lissencephaly patients dominantly disrupt neuronal migration and impair dynein activity. Hum Mol Genet 28:1227–1243. doi:10.1093/hmg/ddy416

Al-Bassam J, Kim H, Flor-Parra I, Lal N, Velji H, Chang F. 2012. Fission yeast Alp14 is a dose-dependent plus end-tracking microtubule polymerase. Mol Biol Cell 23:2878–2890. doi:10.1091/mbc.E12-03-0205

Ayaz P, Munyoki S, Geyer EA, Piedra F-A, Vu ES, Bromberg R, Otwinowski Z, Grishin NV, Brautigam CA, Rice LM. 2014. A tethered delivery mechanism explains the catalytic action of a microtubule polymerase. Elife 3:e03069–e03069. doi:10.7554/eLife.03069

Ayaz P, Ye X, Huddleston P, Brautigam CA, Rice LM. 2012. A TOG:αβ-tubulin Complex Structure Reveals Conformation-Based Mechanisms for a Microtubule Polymerase. Science 337:857. doi:10.1126/science.1221698

Bahi-Buisson N, Poirier K, Fourniol F, Saillour Y, Valence S, Lebrun N, Hully M, Fallet Bianco C, Boddaert N, Elie C, Lascelles K, Souville I, LIS-Tubulinopathies Consortium, Beldjord C, Chelly J. 2014. The wide spectrum of tubulinopathies: what are the key features for the diagnosis? Brain 137:1676–1700. doi:10.1093/brain/awu082

Barkovich AJ, Guerrini R, Kuzniecky RI, Jackson GD, Dobyns WB. 2012. A developmental and genetic classification for malformations of cortical development: update 2012. Brain 135:1348–1369. doi:10.1093/brain/aws019

Barkovich AJ, Kuzniecky RI, Jackson GD, Guerrini R, Dobyns WB. 2005. A developmental and genetic classification for malformations of cortical development. Neurology 65:1873. doi:10.1212/01.wnl.0000183747.05269.2d

Basnet N, Nedozralova H, Crevenna AH, Bodakuntla S, Schlichthaerle T, Taschner M, Cardone G, Janke C, Jungmann R, Magiera MM, Biertümpfel C, Mizuno N. 2018. Direct induction of microtubule branching by microtubule nucleation factor SSNA1. Nat Cell Biol 20:1172– 1180. doi:10.1038/s41556-018-0199-8

Belvindrah R, Natarajan K, Shabajee P, Bruel-Jungerman E, Bernard J, Goutierre M, Moutkine I, Jaglin XH, Savariradjane M, Irinopoulou T, Poncer J-C, Janke C, Francis F. 2017. Mutation of the α-tubulin Tuba1a leads to straighter microtubules and perturbs neuronal migration. J Cell Biol 216:2443–2461. doi:10.1083/jcb.201607074

Bodakuntla S, Jijumon AS, Villablanca C, Gonzalez-Billault C, Janke C. 2019. Microtubule-Associated Proteins: Structuring the Cytoskeleton. Trends in Cell Biology 29:804–819. doi:10.1016/j.tcb.2019.07.004

Borys F, Joachimiak E, Krawczyk H, Fabczak H. 2020. Intrinsic and Extrinsic Factors Affecting Microtubule Dynamics in Normal and Cancer Cells. Molecules 25:3705. doi:10.3390/molecules25163705

Breton S, Brown D. 1998. Cold-induced microtubule disruption and relocalization of membrane proteins in kidney epithelial cells. J Am Soc Nephrol 9:155. doi:10.1681/ASN.V92155

Brouhard GJ, Rice LM. 2018. Microtubule dynamics: an interplay of biochemistry and mechanics. Nat Rev Mol Cell Biol 19:451–463. doi:10.1038/s41580-018-0009-y

Brouhard GJ, Stear JH, Noetzel TL, Al-Bassam J, Kinoshita K, Harrison SC, Howard J, Hyman AA. 2008. XMAP215 is a processive microtubule polymerase. Cell 132:79–88. doi:10.1016/j.cell.2007.11.043

Buey RM, Díaz JF, Andreu JM. 2006. The Nucleotide Switch of Tubulin and Microtubule Assembly: A Polymerization-Driven Structural Change. Biochemistry 45:5933–5938. doi:10.1021/bi060334m

Buscaglia G, Aiken J, Hoff KJ, Northington KR, Bates EA. 2020a. Tuba1a is uniquely important for axon guidance through midline commissural structures. bioRxiv 2020.05.05.079376. doi:10.1101/2020.05.05.079376

Buscaglia G, Northington KR, Moore JK, Bates EA. 2020b. Reduced TUBA1A tubulin causes defects in trafficking and impaired adult motor behavior. eNeuro ENEURO.0045-20.2020. doi:10.1523/ENEURO.0045-20.2020

Chakraborti S, Natarajan K, Curiel J, Janke C, Liu J. 2016. The emerging role of the tubulin code: From the tubulin molecule to neuronal function and disease. Cytoskeleton 73:521–550. doi:10.1002/cm.21290

Chesta ME, Carbajal A, Arce CA, Bisig CG. 2014. Serum-induced neurite retraction in CAD cells – involvement of an ATP-actin retractile system and the lack of microtubule-associated proteins. The FEBS Journal 281:4767–4778. doi:10.1111/febs.12967

Chrétien D, Fuller SD, Karsenti E. 1995. Structure of growing microtubule ends: two-dimensional sheets close into tubes at variable rates. J Cell Biol 129:1311–1328. doi:10.1083/jcb.129.5.1311

Cooper JA. 2014. Molecules and mechanisms that regulate multipolar migration in the intermediate zone. Front Cell Neurosci 8:386–386. doi:10.3389/fncel.2014.00386

Das A, Dickinson DJ, Wood CC, Goldstein B, Slep KC. 2015. Crescerin uses a TOG domain array to regulate microtubules in the primary cilium. Mol Biol Cell 26:4248–4264. doi:10.1091/mbc.E15-08-0603

Dehmelt L, Nalbant P, Steffen W, Halpain S. 2006. A microtubule-based, dynein-dependent force induces local cell protrusions: Implications for neurite initiation. Brain Cell Biology 35:39–56. doi:10.1007/s11068-006-9001-0

Dent EW, Baas PW. 2014. Microtubules in neurons as information carriers. J Neurochem 129:235–239. doi:10.1111/jnc.12621

Dent EW, Gupton SL, Gertler FB. 2011. The growth cone cytoskeleton in axon outgrowth and guidance. Cold Spring Harb Perspect Biol 3:a001800. doi:10.1101/cshperspect.a001800

Dent EW, Kalil K. 2001. Axon branching requires interactions between dynamic microtubules and actin filaments. J Neurosci 21:9757–9769. doi:10.1523/JNEUROSCI.21-24-09757.2001

Dotti CG, Sullivan CA, Banker GA. 1988. The establishment of polarity by hippocampal neurons in culture. J Neurosci 8:1454–1468. doi:10.1523/JNEUROSCI.08-04-01454.1988

Fallet-Bianco C, Laquerrière A, Poirier K, Razavi F, Guimiot F, Dias P, Loeuillet L, Lascelles K, Beldjord C, Carion N, Toussaint A, Revencu N, Addor M-C, Lhermitte B, Gonzales M, Martinovich J, Bessieres B, Marcy-Bonnière M, Jossic F, Marcorelles P, Loget P, Chelly J, Bahi-Buisson N. 2014. Mutations in tubulin genes are frequent causes of various foetal malformations of cortical development including microlissencephaly. Acta Neuropathol Commun 2:69–69. doi:10.1186/2051-5960-2-69

Falnikar A, Tole S, Liu M, Liu JS, Baas PW. 2013. Polarity in migrating neurons is related to a mechanism analogous to cytokinesis. Curr Biol 23:1215–1220. doi:10.1016/j.cub.2013.05.027

Farmer V, Arpağ G, Hall SL, Zanic M. 2021. XMAP215 promotes microtubule catastrophe by disrupting the growing microtubule end. Journal of Cell Biology 220. doi:10.1083/jcb.202012144

Fees CP, Aiken J, O’Toole ET, Giddings TH Jr, Moore JK. 2016. The negatively charged carboxy-terminal tail of β-tubulin promotes proper chromosome segregation. Mol Biol Cell 27:1786–1796. doi:10.1091/mbc.E15-05-0300

Fees CP, Moore JK. 2018. Regulation of microtubule dynamic instability by the carboxy-terminal tail of β-tubulin. Life Sci Alliance 1:e201800054. doi:10.26508/lsa.201800054

Fourniol FJ, Sindelar CV, Amigues B, Clare DK, Thomas G, Perderiset M, Francis F, Houdusse A, Moores CA. 2010. Template-free 13-protofilament microtubule-MAP assembly visualized at 8 A resolution. J Cell Biol 191:463–470. doi:10.1083/jcb.201007081

Gartz Hanson M, Aiken J, Sietsema DV, Sept D, Bates EA, Niswander L, Moore JK. 2016. Novel α-tubulin mutation disrupts neural development and tubulin proteostasis. Dev Biol 409:406–419. doi:10.1016/j.ydbio.2015.11.022

Geyer EA, Burns A, Lalonde BA, Ye X, Piedra F-A, Huffaker TC, Rice LM. 2015. A mutation uncouples the tubulin conformational and GTPase cycles, revealing allosteric control of microtubule dynamics. eLife 4:e10113. doi:10.7554/eLife.10113

Geyer EA, Miller MP, Brautigam CA, Biggins S, Rice LM. 2018. Design principles of a microtubule polymerase. Elife 7:e34574. doi:10.7554/eLife.34574

Gloster A, El-Bizri H, Bamji SX, Rogers D, Miller FD. 1999. Early induction of Tα1 α-tubulin transcription in neurons of the developing nervous system. Journal of Comparative Neurology 405:45–60. doi:10.1002/(SICI)1096-9861(19990301)405:1<45::AID-CNE4>3.0.CO;2-M

Gloster A, Wu W, Speelman A, Weiss S, Causing C, Pozniak C, Reynolds B, Chang E, Toma JG, Miller FD. 1994. The T alpha 1 alpha-tubulin promoter specifies gene expression as a function of neuronal growth and regeneration in transgenic mice. J Neurosci 14:7319– 7330. doi:10.1523/JNEUROSCI.14-12-07319.1994

Goodson HV, Jonasson EM. 2018. Microtubules and Microtubule-Associated Proteins. Cold Spring Harb Perspect Biol 10:a022608. doi:10.1101/cshperspect.a022608

Hahn I, Voelzmann A, Parkin J, Fülle JB, Slater PG, Lowery LA, Sanchez-Soriano N, Prokop A. 2021. Tau, XMAP215/Msps and Eb1 co-operate interdependently to regulate microtubule polymerisation and bundle formation in axons. PLOS Genetics 17:e1009647. doi:10.1371/journal.pgen.1009647

Hand R, Bortone D, Mattar P, Nguyen L, Heng JI-T, Guerrier S, Boutt E, Peters E, Barnes AP, Parras C, Schuurmans C, Guillemot F, Polleux F. 2005. Phosphorylation of Neurogenin2 Specifies the Migration Properties and the Dendritic Morphology of Pyramidal Neurons in the Neocortex. Neuron 48:45–62. doi:10.1016/j.neuron.2005.08.032

He Y, Yu W, Baas PW. 2002. Microtubule Reconfiguration during Axonal Retraction Induced by Nitric Oxide. J Neurosci 22:5982. doi:10.1523/JNEUROSCI.22-14-05982.2002

Hebebrand M, Hüffmeier U, Trollmann R, Hehr U, Uebe S, Ekici AB, Kraus C, Krumbiegel M, Reis A, Thiel CT, Popp B. 2019. The mutational and phenotypic spectrum of TUBA1A-associated tubulinopathy. Orphanet J Rare Dis 14:38–38. doi:10.1186/s13023-019-1020-x

Horio T, Hotani H. 1986. Visualization of the dynamic instability of individual microtubules by dark-field microscopy. Nature 321:605–607. doi:10.1038/321605a0

Hussmann F, Drummond DR, Peet DR, Martin DS, Cross RA. 2016. Alp7/TACC-Alp14/TOG generates long-lived, fast-growing MTs by an unconventional mechanism. Sci Rep 6:20653–20653. doi:10.1038/srep20653

Jánosi IM, Chrétien D, Flyvbjerg H. 1998. Modeling elastic properties of microtubule tips and walls. European Biophysics Journal 27:501–513. doi:10.1007/s002490050160

Jordan MA, Toso RJ, Thrower D, Wilson L. 1993. Mechanism of mitotic block and inhibition of cell proliferation by taxol at low concentrations. Proc Natl Acad Sci U S A 90:9552–9556. doi:10.1073/pnas.90.20.9552

Kapitein LC, Hoogenraad CC. 2015. Building the Neuronal Microtubule Cytoskeleton. Neuron 87:492–506. doi:10.1016/j.neuron.2015.05.046

Keays DA, Tian G, Poirier K, Huang G-J, Siebold C, Cleak J, Oliver PL, Fray M, Harvey RJ, Molnár Z, Piñon MC, Dear N, Valdar W, Brown SDM, Davies KE, Rawlins JNP, Cowan NJ, Nolan P, Chelly J, Flint J. 2007. Mutations in alpha-tubulin cause abnormal neuronal migration in mice and lissencephaly in humans. Cell 128:45–57. doi:10.1016/j.cell.2006.12.017

Kosco KA, Pearson CG, Maddox PS, Wang PJ, Adams IR, Salmon ED, Bloom K, Huffaker TC. 2001. Control of microtubule dynamics by Stu2p is essential for spindle orientation and metaphase chromosome alignment in yeast. Mol Biol Cell 12:2870–2880. doi:10.1091/mbc.12.9.2870

Lasser M, Tiber J, Lowery LA. 2018. The Role of the Microtubule Cytoskeleton in Neurodevelopmental Disorders. Frontiers in Cellular Neuroscience 12:165. doi:10.3389/fncel.2018.00165

Lee MJ, Gergely F, Jeffers K, Peak-Chew SY, Raff JW. 2001. Msps/XMAP215 interacts with the centrosomal protein D-TACC to regulate microtubule behaviour. Nature Cell Biology 3:643–649. doi:10.1038/35083033

Lin S, Liu M, Mozgova OI, Yu W, Baas PW. 2012. Mitotic motors coregulate microtubule patterns in axons and dendrites. J Neurosci 32:14033–14049. doi:10.1523/JNEUROSCI.3070-12.2012

Löwe J, Li H, Downing KH, Nogales E. 2001. Refined structure of αβ-tubulin at 3.5 Å resolution11Edited by I. A. Wilson. Journal of Molecular Biology 313:1045–1057. doi:10.1006/jmbi.2001.5077

Lowery LA, Stout A, Faris AE, Ding L, Baird MA, Davidson MW, Danuser G, Van Vactor D. 2013. Growth cone-specific functions of XMAP215 in restricting microtubule dynamics and promoting axonal outgrowth. Neural Dev 8:22–22. doi:10.1186/1749-8104-8-22

Lu W, Fox P, Lakonishok M, Davidson MW, Gelfand VI. 2013. Initial Neurite Outgrowth in Drosophila Neurons Is Driven by Kinesin-Powered Microtubule Sliding. Current Biology 23:1018–1023. doi:10.1016/j.cub.2013.04.050

Ludueña RF, Banerjee A. 2008. The Tubulin Superfamily In: Fojo T, editor. The Role of Microtubules in Cell Biology, Neurobiology, and Oncology. Totowa, NJ: Humana Press. pp. 177–191. doi:10.1007/978-1-59745-336-3_7

Mahamdeh M, Simmert S, Luchniak A, Schäffer E, Howard J. 2018. Label-free high-speed wide-field imaging of single microtubules using interference reflection microscopy. Journal of Microscopy 272:60–66. doi:10.1111/jmi.12744

Mandelkow EM, Mandelkow E, Milligan RA. 1991. Microtubule dynamics and microtubule caps: a time-resolved cryo-electron microscopy study. Journal of Cell Biology 114:977–991. doi:10.1083/jcb.114.5.977

Manka SW, Moores CA. 2020. Pseudo-repeats in doublecortin make distinct mechanistic contributions to microtubule regulation. EMBO Rep 21:e51534–e51534. doi:10.15252/embr.202051534

Manka SW, Moores CA. 2018. Microtubule structure by cryo-EM: snapshots of dynamic instability. Essays Biochem 62:737–751. doi:10.1042/EBC20180031

Matsuo Y, Maurer SP, Yukawa M, Zakian S, Singleton MR, Surrey T, Toda T. 2016. An unconventional interaction between Dis1/TOG and Mal3/EB1 in fission yeast promotes the fidelity of chromosome segregation. J Cell Sci 129:4592–4606. doi:10.1242/jcs.197533

Meijering E, Jacob M, Sarria J-CF, Steiner P, Hirling H, Unser M. 2004. Design and validation of a tool for neurite tracing and analysis in fluorescence microscopy images. Cytometry Part A 58A:167–176. doi:10.1002/cyto.a.20022

Miller RK. 2004. Monitoring Spindle Assembly and Disassembly in Yeast by Indirect Immunofluorescence In: Lieberman HB, editor. Cell Cycle Checkpoint Control Protocols. Totowa, NJ: Humana Press. pp. 341–352. doi:10.1385/1-59259-646-0:341

Minoura I, Hachikubo Y, Yamakita Y, Takazaki H, Ayukawa R, Uchimura S, Muto E. 2013. Overexpression, purification, and functional analysis of recombinant human tubulin dimer. FEBS Letters 587:3450–3455. doi:10.1016/j.febslet.2013.08.032

Mitchison T, Kirschner M. 1984. Dynamic instability of microtubule growth. Nature 312:237–242. doi:10.1038/312237a0

Moores CA, Perderiset M, Kappeler C, Kain S, Drummond D, Perkins SJ, Chelly J, Cross R, Houdusse A, Francis F. 2006. Distinct roles of doublecortin modulating the microtubule cytoskeleton. EMBO J 25:4448–4457. doi:10.1038/sj.emboj.7601335

Nadarajah B, Brunstrom JE, Grutzendler J, Wong ROL, Pearlman AL. 2001. Two modes of radial migration in early development of the cerebral cortex. Nature Neuroscience 4:143–150. doi:10.1038/83967

Nawrotek A, Knossow M, Gigant B. 2011. The Determinants That Govern Microtubule Assembly from the Atomic Structure of GTP-Tubulin. Journal of Molecular Biology 412:35–42. doi:10.1016/j.jmb.2011.07.029

Noctor SC, Martínez-Cerdeño V, Ivic L, Kriegstein AR. 2004. Cortical neurons arise in symmetric and asymmetric division zones and migrate through specific phases. Nature Neuroscience 7:136–144. doi:10.1038/nn1172

Nogales E, Wang H-W. 2006. Structural mechanisms underlying nucleotide-dependent self-assembly of tubulin and its relatives. Current Opinion in Structural Biology 16:221–229. doi:10.1016/j.sbi.2006.03.005

Ohshima T, Hirasawa M, Tabata H, Mutoh T, Adachi T, Suzuki H, Saruta K, Iwasato T, Itohara S, Hashimoto M, Nakajima K, Ogawa M, Kulkarni AB, Mikoshiba K. 2007. Cdk5 is required for multipolar-to-bipolar transition during radial neuronal migration and proper dendrite development of pyramidal neurons in the cerebral cortex. Development 134:2273–2282. doi:10.1242/dev.02854

Podolski M, Mahamdeh M, Howard J. 2014. Stu2, the budding yeast XMAP215/Dis1 homolog, promotes assembly of yeast microtubules by increasing growth rate and decreasing catastrophe frequency. J Biol Chem 289:28087–28093. doi:10.1074/jbc.M114.584300

Ravelli RBG, Gigant B, Curmi PA, Jourdain I, Lachkar S, Sobel A, Knossow M. 2004. Insight into tubulin regulation from a complex with colchicine and a stathmin-like domain. Nature 428:198–202. doi:10.1038/nature02393

Reusch S, Biswas A, Hirst WG, Reber S. 2020. Affinity Purification of Label-free Tubulins from Xenopus Egg Extracts. STAR Protoc 1:100151–100151. doi:10.1016/j.xpro.2020.100151

Rice LM, Montabana EA, Agard DA. 2008. The lattice as allosteric effector: structural studies of alphabeta- and gamma-tubulin clarify the role of GTP in microtubule assembly. Proc Natl Acad Sci U S A 105:5378–5383. doi:10.1073/pnas.0801155105

Roostalu J, Thomas C, Cade NI, Kunzelmann S, Taylor IA, Surrey T. 2020. The speed of GTP hydrolysis determines GTP cap size and controls microtubule stability. eLife 9:e51992. doi:10.7554/eLife.51992

Sainath R, Gallo G. 2015. Cytoskeletal and signaling mechanisms of neurite formation. Cell and Tissue Research 359:267–278. doi:10.1007/s00441-014-1955-0

Slater PG, Cammarata GM, Samuelson AG, Magee A, Hu Y, Lowery LA. 2019. XMAP215 promotes microtubule-F-actin interactions to regulate growth cone microtubules during axon guidance in Xenopus laevis. J Cell Sci 132:jcs224311. doi:10.1242/jcs.224311

Slep KC, Vale RD. 2007. Structural basis of microtubule plus end tracking by XMAP215, CLIP-170, and EB1. Mol Cell 27:976–991. doi:10.1016/j.molcel.2007.07.023

Srivastava AK, Schwartz CE. 2014. Intellectual disability and autism spectrum disorders: Causal genes and molecular mechanisms. Neuroscience & Biobehavioral Reviews 46:161–174. doi:10.1016/j.neubiorev.2014.02.015

Studier FW. 2005. Protein production by auto-induction in high-density shaking cultures. Protein Expression and Purification 41:207–234. doi:10.1016/j.pep.2005.01.016

Tabata H, Nakajima K. 2003. Multipolar Migration: The Third Mode of Radial Neuronal Migration in the Developing Cerebral Cortex. J Neurosci 23:9996. doi:10.1523/JNEUROSCI.23-31-09996.2003

Tilney LG, Porter KR. 1967. Studies on the microtubules in heliozoa. II. The effect of low temperature on these structures in the formation and maintenance of the axopodia. J Cell Biol 34:327–343. doi:10.1083/jcb.34.1.327

van der Vaart B, Manatschal C, Grigoriev I, Olieric V, Gouveia SM, Bjelic S, Demmers J, Vorobjev I, Hoogenraad CC, Steinmetz MO, Akhmanova A. 2011. SLAIN2 links microtubule plus end-tracking proteins and controls microtubule growth in interphase. J Cell Biol 193:1083–1099. doi:10.1083/jcb.201012179

Vasquez RJ, Gard DL, Cassimeris L. 1994. XMAP from Xenopus eggs promotes rapid plus end assembly of microtubules and rapid microtubule polymer turnover. J Cell Biol 127:985– 993. doi:10.1083/jcb.127.4.985

Walker RA, O’Brien ET, Pryer NK, Soboeiro MF, Voter WA, Erickson HP, Salmon ED. 1988. Dynamic instability of individual microtubules analyzed by video light microscopy: rate constants and transition frequencies. J Cell Biol 107:1437–1448. doi:10.1083/jcb.107.4.1437

Weber K, Pollack R, Bibring T. 1975. Antibody against tuberlin: the specific visualization of cytoplasmic microtubules in tissue culture cells. Proc Natl Acad Sci U S A 72:459–463. doi:10.1073/pnas.72.2.459

Westermann S, Weber K. 2003. Post-translational modifications regulate microtubule function. Nature Reviews Molecular Cell Biology 4:938–948. doi:10.1038/nrm1260

Widlund PO, Podolski M, Reber S, Alper J, Storch M, Hyman AA, Howard J, Drechsel DN. 2012. One-step purification of assembly-competent tubulin from diverse eukaryotic sources. Mol Biol Cell 23:4393–4401. doi:10.1091/mbc.E12-06-0444

Winding M, Kelliher MT, Lu W, Wildonger J, Gelfand VI. 2016. Role of kinesin-1–based microtubule sliding in *Drosophila* nervous system development. Proc Natl Acad Sci USA 113:E4985. doi:10.1073/pnas.1522416113

Witte H, Neukirchen D, Bradke F. 2008. Microtubule stabilization specifies initial neuronal polarization. J Cell Biol 180:619–632. doi:10.1083/jcb.200707042

Zanic M, Widlund PO, Hyman AA, Howard J. 2013. Synergy between XMAP215 and EB1 increases microtubule growth rates to physiological levels. Nature Cell Biology 15:688– 693. doi:10.1038/ncb2744

Zheng Y, Wong ML, Alberts B, Mitchison T. 1995. Nucleation of microtubule assembly by a γ-tubulin-containing ring complex. Nature 378:578–583. doi:10.1038/378578a0

